# Structural basis of selective phospholipid recognition and local membrane deformation by CTL1

**DOI:** 10.64898/2026.10.02.755950

**Authors:** Hanxing Jiang, Adrián Martínez-Jiménez, Kazi Amirul Hossain, Haigang Song, Borja Ochoa-Lizarralde, Jorge Pedro López-Alonso, Guillem Pont-Espinós, Raúl Estevez, Modesto Orozco, Carol V. Robinson, Iban Ubarretxena-Belandia, Igor Tascón, Ekaitz Errasti-Murugarren

**Affiliations:** Instituto Biofisika (UPV/EHU, CSIC), Leioa, Spain; Physiological Sciences Department, IDIBELL-Institute of Neurosciences, School of Medicine and Health Sciences, University of Barcelona, Bellvitge Campus. L’Hospitalet de Llobregat, Spain; Institute for Research in Biomedicine (IRB Barcelona), The Barcelona Institute of Science and Technology (BIST), Baldiri Reixac 10, E-08028, Barcelona, Spain; Kavli Institute for Nanoscience Discovery, Dorothy Crowfoot Hodgkin Building, University of Oxford, OX1 3QU, United Kingdom; Department of Chemistry, University of Oxford, OX1 3TA, United Kingdom; Basque Resource for Electron Microscopy, Leioa, Spain; Center for Biomedical Network Research of Rare Diseases (CIBERER U-731), ISCIII, Madrid, Spain; Ikerbasque Foundation for Science, Bilbao, Spain

**Author notes:** Corresponding authors (I.U-B), (I.T), (E.E-M).

**Keywords:** SLC44A1, cryo-electron microscopy, phospholipid-binding protein, membrane remodelling

## Abstract

The physiological function of the human choline transporter-like protein CTL1 (SLC44A1) remains elusive despite its implication in key pathophysiological processes. Here we combine cryo-electron microscopy, native mass spectrometry, lipidomics, molecular dynamics simulations and functional assays to characterise the molecular basis of phospholipid recognition by CTL1 and explore its potential molecular role. High-resolution structures of CTL1 in both monomeric and dimeric states reveal a broad amphipathic cavity and an extracellular domain that accommodate multiple phospholipid species, identifying phosphatidylethanolamine, phosphatidylserine, phosphatidylinositol and phosphatidic acid as endogenous CTL1 ligands. Molecular dynamics simulations show spontaneous phospholipid recruitment into these conserved binding sites accompanied by membrane thinning, curvature and local phospholipid reorganisation, with membrane deformation being enhanced upon dimerisation. Functional analyses in mammalian cells and in reconstituted proteoliposomes failed to detect CTL1-mediated choline transport, supporting a function distinct from canonical choline uptake. Together, our findings suggest that CTL1 acts as a phospholipid-binding protein capable of locally deforming the membrane and provide a structural framework linking phospholipid recognition to membrane organisation. These results offer new insights into potential mechanisms underlying the altered phospholipid homeostasis associated with CTL1 dysfunction in neurological diseases and cancer.

## Introduction

Cell membranes are highly organised structures, and their lipid composition determines not only their physical properties but also the localisation, trafficking and activity of membrane proteins^1^. Rather than serving as passive barriers, membranes are dynamic assemblies in which membrane proteins and surrounding phospholipids reciprocally influence one another to regulate membrane organisation and protein localisation^2–5^. Although several membrane proteins have been shown to alter local membrane architecture^1–4^, the molecular mechanisms by which they regulate phospholipid organisation remain poorly understood. Understanding these mechanisms is therefore essential for explaining how membrane architecture is established and how its disruption contributes to disease.

CTL1 (SLC44A1) is expressed in essentially all human cell types and thought to localise primarily to the plasma membrane, yet its physiological role remains unresolved despite over two decades of investigation. Initially described as a Na -independent choline transporter^6–10^, CTL1 has subsequently been implicated in phospholipid metabolism^11^, membrane trafficking^12–14^, myelination^15–17^, and neurological function ^18–23^. Despite these observations, the molecular activity of CTL1 and the mechanism linking CTL1 dysfunction to phospholipid homeostasis and membrane protein organisation remain poorly understood.

Here, we combine cryo-electron microscopy (cryo-EM), functional assays, native mass spectrometry (nMS), lipidomics, proteomics, and molecular dynamics (MD) simulations to investigate the molecular function of human CTL1. We show that CTL1 adopts the recently described SLC44 fold^24,25^ in both monomeric and dimeric states, forming a broad amphipathic cavity that extends from the membrane surface towards the protein core. We identify multiple conserved phospholipid-binding sites and establish phosphatidylethanolamine (PE), phosphatidylserine (PS), phosphatidylinositol (PI) and phosphatidic acid (PA) as endogenous CTL1 ligands. MD simulations further reveal spontaneous phospholipid recruitment into these sites, accompanied by local membrane thinning, curvature and phospholipid reorganisation, suggesting a close relationship between phospholipid binding and local membrane deformation.

Together, these findings support a functional reinterpretation of CTL1 as a phospholipid-binding membrane protein associated with local membrane deformation, rather than as a conventional choline transporter. They provide a molecular framework for investigating how CTL1 may contribute to phospholipid homeostasis and how its dysfunction may be linked to neurodegeneration, myelination defects, and cancer-associated metabolic adaptation^26–32^.

## Results

### Reassessment of CTL1-mediated choline uptake

CTL1 has long been proposed to function as a Na^+^-independent choline transporter. To determine whether increased CTL1 expression is sufficient to enhance choline uptake, we transiently overexpressed human CTL1 in HCT116 cells and measured [^3^H]-choline uptake under experimental conditions previously used to characterise CTL1-mediated choline transport in multiple mammalian cell lines^7,33^, including choline concentrations ranging from 1 to 100 μM, Na^+^-containing or Na^+^-free (NMDG) transport buffers, and extracellular pH values of 5.5 and 7.5 (**Fig. 1a**). Across all experimental conditions tested, CTL1 overexpression did not produce a measurable increase in choline uptake compared with cells transfected with the empty vector, despite strong accumulation of CTL1 at the plasma membrane (**Fig. 1b**). Choline uptake remained essentially unchanged irrespective of extracellular Na^+^ concentration or pH, indicating that increased plasma membrane expression of CTL1 alone is insufficient to enhance cellular choline transport.

**Figure 1.**
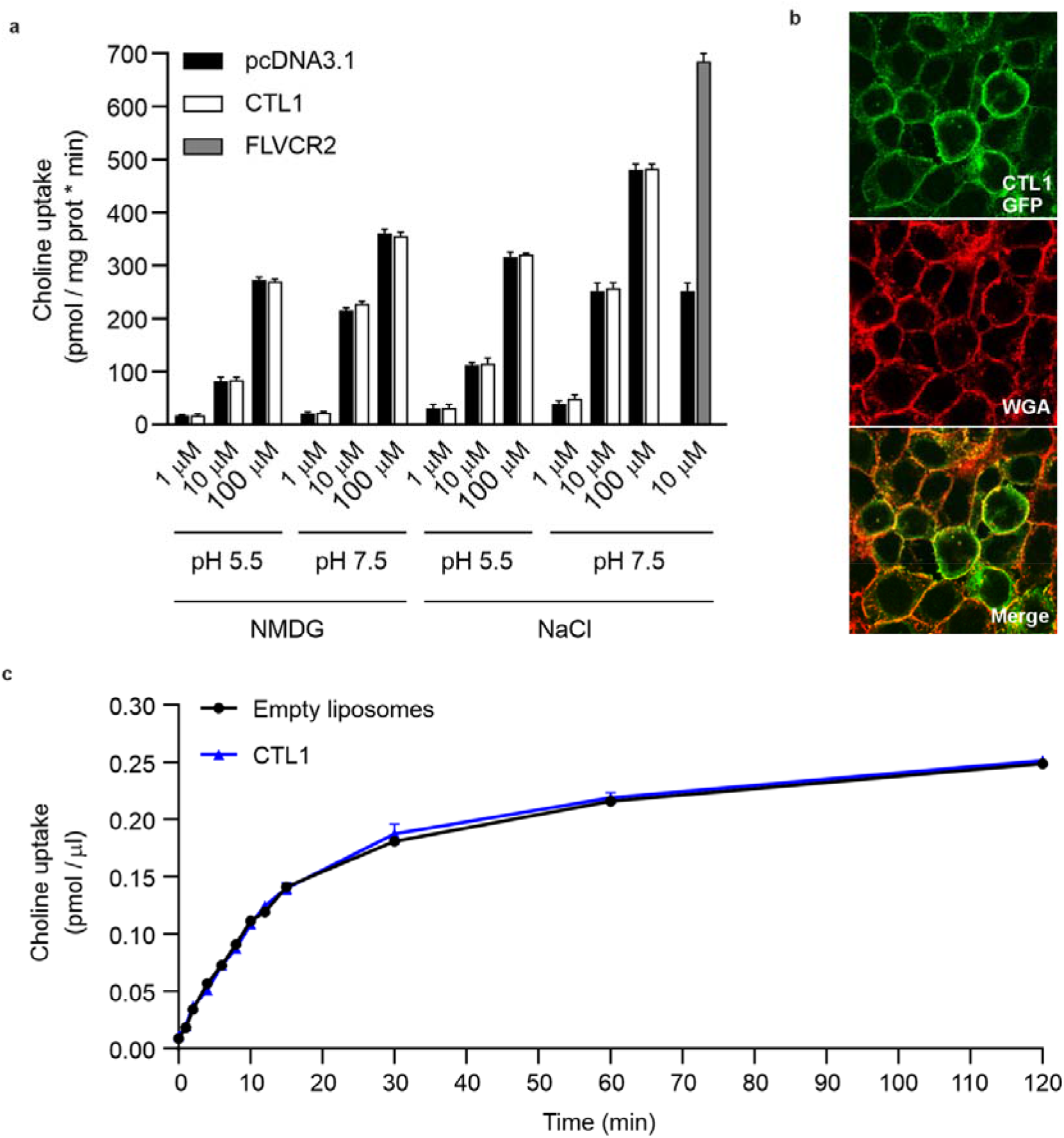
Absence of CTL1-mediated choline uptake in HCT116 cells. **a** Initial rates of [^3^H]-choline uptake (pmol choline/mg protein•min) in HCT116 cells transiently expressing human CTL1 (CTL1-pcDNA3.1), human FLVCR2 (FLVCR2-pcDNA3.1), or empty vector (pcDNA3.1). Uptake was measured after a 1 min incubation over a range of extracellular choline concentrations (1, 10 and 100 μM), in either Na^+^-containing (137 mM NaCl) or Na^+^-free (137 mM NMDG) transport buffers, and at extracellular pH 7.5 or 5.5. Across all conditions tested, CTL1 overexpression did not increase choline uptake above control cells, whereas overexpression of FLVCR2 correlated with robust choline uptake. Data (mean ± SD) correspond to three independent experiments run in quadruplicate (N=3). **b** Subcellular localisation of GFP-tagged human CTL1 in HCT116 cells. Colour code: CTL1-GFP (green) and Wheat Germ Agglutinin (WGA; red). Merged image shows CTL1 localises at the plasma membrane (identified by WGA) of cells. **c** Transport activity of human CTL1 reconstituted into total brain lipid liposomes. In their lumen, liposomes contained 20 mM Tris-HCl (pH 7.4), 150 mM KCl and 1 mM CaCl_2_. Time course (0 to 120 min) of 50 μM [^3^H]-choline (2 μCi/ml) uptake (pmol choline/µl) into CTL1 proteoliposomes (blue triangles) and into empty liposomes (black circles) in a Na^+^-containing uptake buffer (10 mM HEPES pH 7.4, 150 mM NaCl, 2 mM CaCl_2_, and 1 mM MgSO_4_). Data correspond to a representative experiment run in triplicate (N=3). A second independent experiment gave similar results.

To exclude the possibility that this observation was specific to HCT116 cells, we extended the experiments to additional mammalian cell lines, including HeLa, HEK293T, CHO and MDCK cells (**Supplementary Fig. 1a**). Consistent with the results obtained in HCT116 cells, transient CTL1 overexpression failed to increase choline uptake in any cellular background examined, despite efficient plasma membrane localisation of CTL1 in all cell lines, as confirmed by fluorescence microscopy (**Supplementary Fig. 1b**). As a positive control, parallel experiments were performed using FLVCR2, a recently established high-affinity choline transporter^34,35^. In contrast to CTL1, FLVCR2 overexpression produced a marked increase in choline uptake in all cell lines analysed, confirming that the assay conditions were fully capable of detecting transporter-dependent choline uptake (**Fig. 1a and Supplementary Fig. 1a**).

To rule out whether the cellular environment had somehow masked CTL1-mediated transport activity, we reconstituted detergent-purified CTL1 into liposomes prepared from porcine brain total lipid extracts (**Supplementary Fig. 2a-c**). Successful incorporation of CTL1 into proteoliposomes was confirmed by immunoblotting (**Supplementary Fig. 2c**), after which [^3^H]-choline uptake was measured using our previously validated proteoliposome assay^36^ under the same experimental conditions employed in the cellular transport assays. Consistent with the results obtained in cells, CTL1-containing proteoliposomes showed negligible choline uptake above empty-liposome controls throughout the time course examined (**Fig. 1c**). The absence of measurable transport activity in both cellular and reconstituted systems prompted us to investigate the molecular function of CTL1 through structural and biophysical approaches.

### High-resolution structures of human CTL1 reveal the previously unresolved extracellular domain and defines interactions with endogenous lipids

To investigate the structural basis of CTL1 function, full-length human CTL1 was expressed in HEK293F cells and purified in mixed micelles of n-dodecyl-β-D-maltoside (DDM) and cholesterol hemisuccinate (CHS). Size-exclusion chromatography (SEC) resolved monomeric and dimeric species (**Supplementary Fig. 3a, b**), and cryo-EM study yielded high-resolution 3D reconstructions for both at a nominal resolution of 3.05 Å and 3.23 Å, respectively (**Figs. 2a, 3a**, **Supplementary Fig. 3c, d, and Table 1**). Cryo-EM maps enabled near-complete *de novo* model building of CTL1, including the first structural determination of the extensive extracellular domain (ECD; residues 53-211) (**Fig. 2b**, **Supplementary Fig. 4**). Only residues 1-15 at the N-terminus and 629-657 at the C-terminus remained unresolved.

**Figure 2.**
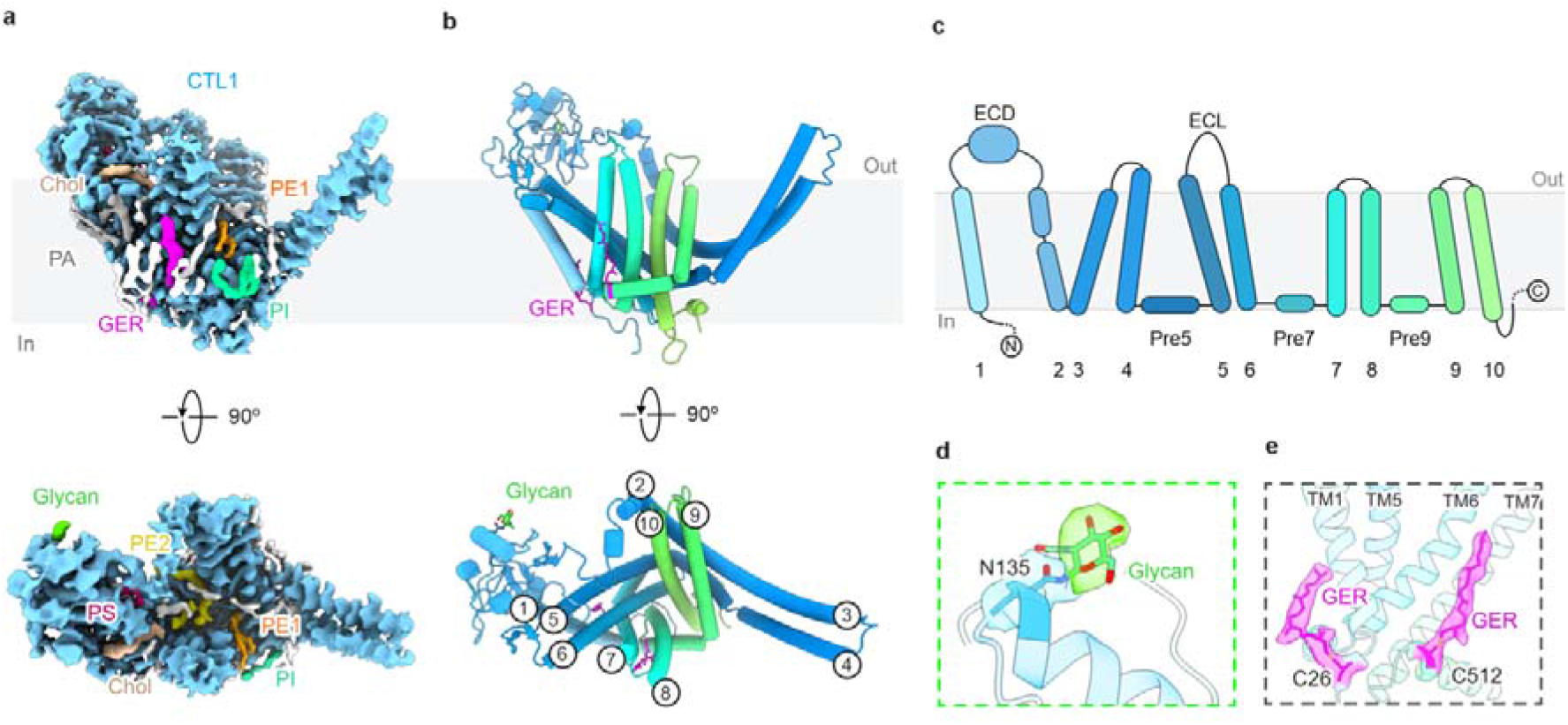
Structure of monomeric human CTL1. **a** Cryo-EM map of the CTL1 monomer shown in blue, with bound lipids and post-translational modifications coloured individually, and unassigned densities shown in white. Grey slab represents the membrane. **b** Cartoon representation of CTL1 monomer atomic model, coloured from the N-terminus (blue) to the C-terminus (green). Geranylgeranylation (GER; magenta) and the N-linked glycan (lime green) are shown as sticks. Transmembrane (TM) helices are numbered in the top view (bottom panel). **c** Topology diagram of CTL1 monomer. Dashed lines indicate unresolved N-terminal and C-terminal extensions. **d** Close-up view of the N-linked glycan (lime green) and N135 (blue) shown as sticks within the cryo-EM density. **e** Close-up view of the two geranylgeranylated cysteines (C26 and C512) shown as magenta sticks within their respective cryo-EM densities.

**Figure 3.**
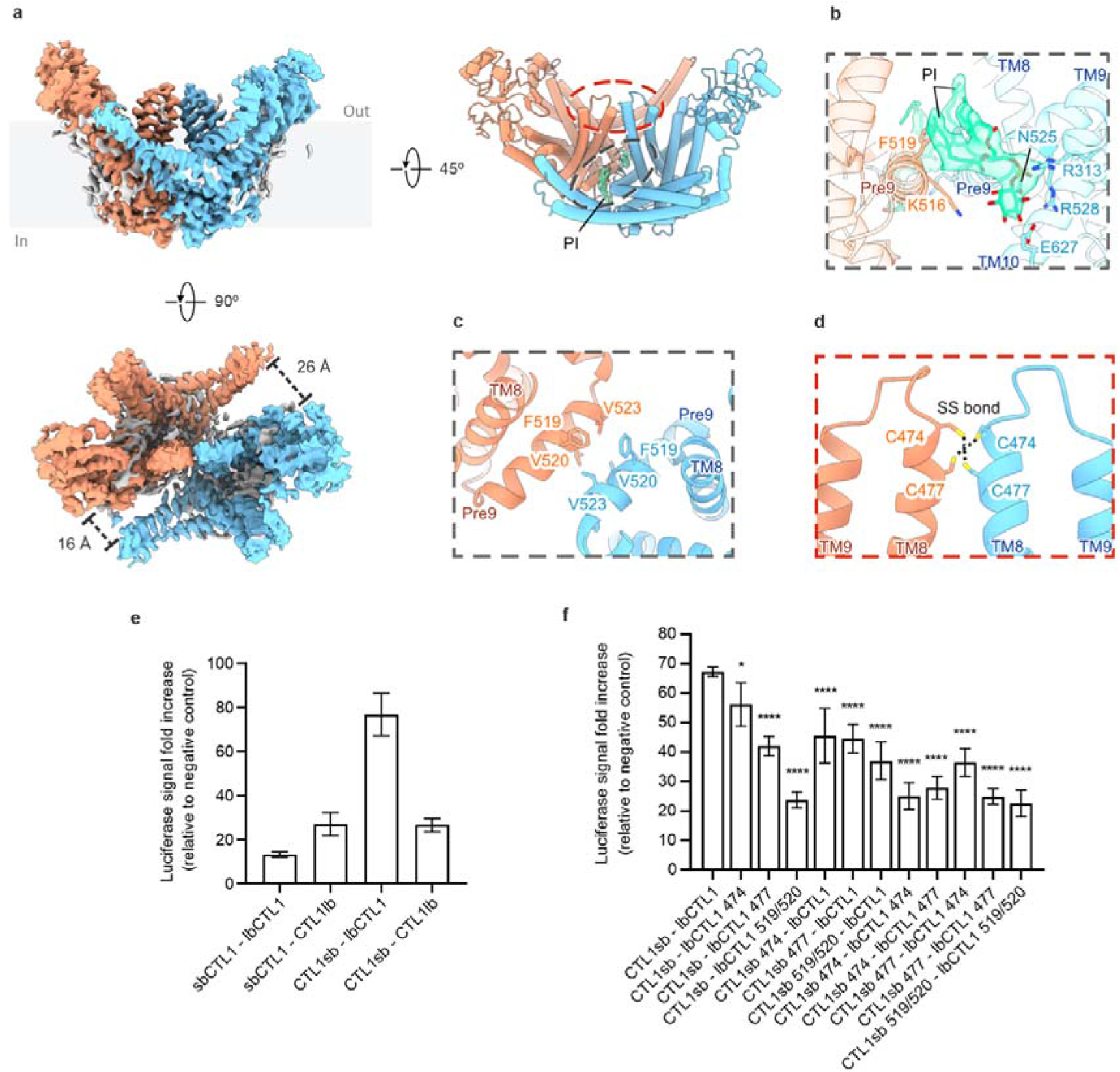
Structural overview and characterisation of CTL1 dimerisation. **a** Cryo-EM map of CTL1 homodimer with protomers depicted in blue and salmon, and surrounding lipid densities in white. Right, 45°-rotated cartoon representation highlighting the dimerisation interface formed by TM8 (red dashed line) and Pre9 (black dashed line) of each protomer. Two PI molecules are shown as sticks within the cryo-EM densities (light green). Bottom, top view of the dimer with the distances TM3/TM4-ECD in both protomers. **b** Close-up view as outlined in **a** (black dashed line) shows interactions between a PI molecule (light green) and residues from each protomer. **c** Close-up view of the hydrophobic contacts between Pre9 residues from both protomers as indicated in **a** (black dashed line). **d** Close-up view as outlined in **a** (red dashed line) shows intermolecular disulfide bonds (dotted lines) formed between TM8 cysteine residues from both protomers. **e** Identification of the optimal NanoBiT probe pairs for detecting CTL1 dimerisation. Pair names indicate the N- or C-terminal orientation of LgBiT (lb) and SmBiT (sb). Data (mean ± SD) correspond to six independent experiments run in triplicate (N=6). **f** Effect of C474S, C477S, F519A and V520A substitutions on CTL1 dimerisation measured by NanoBiT complementation. Luciferase fold increase was calculated relative to the corresponding negative control. Data (mean ± SD) correspond to at least three independent experiments run in triplicate (N=3-6). Statistical significance was determined by two-way ANOVA followed by Dunnett’s multiple-comparisons test using the WT-WT condition as the control. ∗∗∗∗ *p* < 0.0001, F(11,41) = 35.97.

**Table 1.**
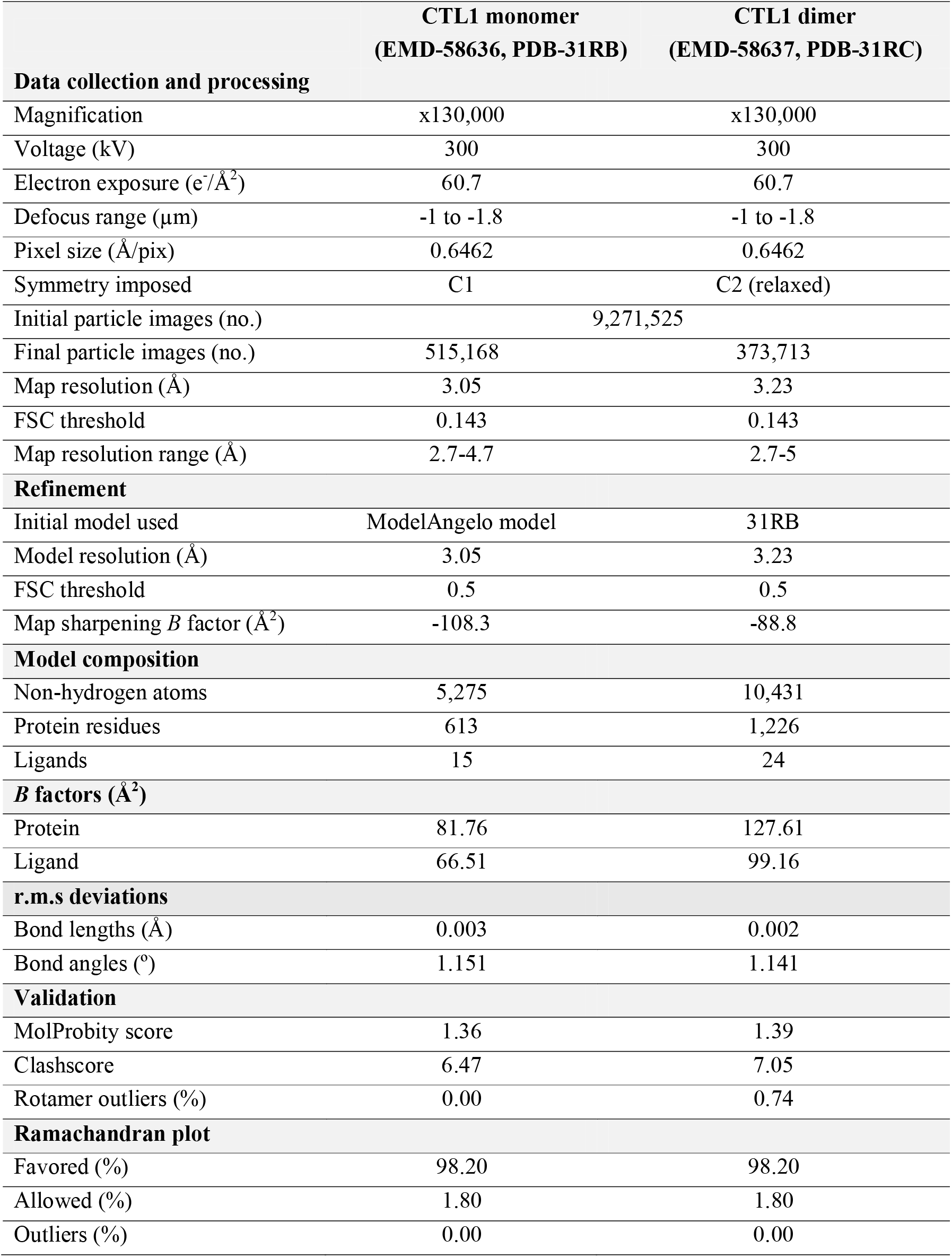
Cryo-EM structure determination. Cryo-EM data collection, refinement and validation statistics.

In agreement with previous studies^24,25^, CTL1 consists of ten transmembrane (TM) helices (TM1-TM10) with intracellular N- and C-termini and a large ECD inserted between TM1 and TM2 (**Fig. 2b, c**). The TM domains display a pseudo-radial arrangement, with three consecutive hairpins formed by TM5-TM10 that open toward the extracellular side forming the structural core around a central cavity, whereas TM1-TM4 occupy peripheral positions. At the intracellular leaflet, three short amphipathic helices parallel to the membrane (Pre5, Pre7 and Pre9) connect TM4-TM5, TM6-TM7 and TM8-TM9, respectively (**Fig. 2c**). The ECD comprises short helices and β-sheets stabilised by four pairs of disulfide bonds (**Supplementary Fig. 5a**). In addition, an extracellular loop (ECL), linking TM5 and TM6, is attached to the ECD through cation-π interaction between R57 and F375, as well as hydrophobic contacts between V110 and F372, likely contributing to local structural stability (**Supplementary Fig. 5a**). The ECD adopts a fold that, to our knowledge, has not been previously described and appears to be unique to CTL1 among SLC44 family members (**Supplementary Fig. 5b, c and Supplementary Fig. 6**).

nMS supported by proteomics revealed a diverse set of post-translational modifications (PTMs), including N-myristoylation, geranylgeranylation, palmitoylation and heterogeneous high-mannose N-glycosylation **(Supplementary Fig. 7a-c)**. Inspection of the cryo-EM structure revealed a glycan-like density attached to N135 in the ECD (**Fig. 2d**). Consistent with this assignment, glycosylated species detected by nMS were absent in the CTL1-N135A variant (**Supplementary Fig. 7c**), indicating that N135 is the major detectable N-glycosylation site. Additional densities were observed at C26 and C512, consistent with geranylgeranylation of these cysteine residues (**Fig. 2e and Supplementary Fig. 7a, c**). Both modified cysteines are located at the inner leaflet, with their hydrophobic chains oriented toward the bilayer core, likely contributing to CTL1 anchoring and stability within specific membrane regions^37^.

In addition, the cryo-EM maps revealed several strong non-proteinaceous densities surrounding the TM domains (**Supplementary Fig. 4 and 8**). Although CHS present in the solubilisation detergent mixture could potentially account for these densities, cholesterol provides a substantially better fit (**Supplementary Fig. 8b**). This suggests that endogenous cholesterol molecules remained tightly bound to CTL1 throughout purification and were consequently modelled in the structure. Three cholesterol molecules (Chol1-Chol3) are closely arranged along a hydrophobic groove on the membrane-facing surfaces of TM1, TM5, TM6 and the Pre7-TM7 region (**Supplementary Fig. 8a-c**). Within this cluster, the central cholesterol (Chol2), buried between TM5 and TM6, forms a hydrogen bond with Q337, and may promote cooperative stabilisation of the neighbouring cholesterol molecules through van der Waals contacts (**Supplementary Fig. 8c**). An elongated lipid-like density resembling an acyl chain extends through the groove over Chol2 and engages in hydrophobic contacts with the aliphatic tail of Chol3, potentially further stabilising the assembly (**Supplementary Fig. 8c**). The shape of the density is compatible with a lysophosphatidylcholine (LPC) molecule, however its hydrophobic contacts with the neighbouring cholesterol molecule are inconsistent with LPC and instead favour a plasmalogen LPC (P-LPC) (**Supplementary Fig. 8f**). Together with recent evidence implicating CTL1 in plasmalogen biology^16^, these observations led us to tentatively model this density as a P-LPC molecule. In this model, the phosphate group coordinates a Mg^2+^ ion, interacting with the main chain of L333 and side chain of Q337 and W340 (**Supplementary Fig. 8d**). Additionally, a fourth cholesterol molecule (Chol4) was found near the extracellular boundary of the membrane, where it is held within a hydrophobic pocket formed by residues from TM5 and TM6 (**Supplementary Fig. 8a, e**). This cholesterol is further stabilised by a cation-π interaction of its tetracyclic ring with R105, and a hydrogen bond between its hydroxyl group and K376 (**Supplementary Fig. 8e**). Remarkably, two additional cryo-EM densities were observed on the surface of the protein adjacent to Chol4. Based on the characteristic shape of the polar headgroup densities, we assigned these densities as PA lipid molecules (**Supplementary Fig. 8e**). Together, these observations indicate that CTL1 co-purifies with multiple endogenous lipids, consistent with a close association between the protein and its surrounding lipid environment.

### Molecular determinants of CTL1 dimerisation

Dimeric CTL1 accounted for ∼35% of the cryo-EM dataset. The dimer is formed primarily through interactions between TM8 and the Pre9 helix of each protomer, burying ∼463 Å^2^ of surface area (**Fig. 3a**). Two symmetrically positioned phosphatidylinositol (PI) molecules are observed at the dimer interface, likely enhancing dimer contacts. The PI headgroups are coordinated by Pre5 R313, Pre9 N525 and R528, and C-terminal E627, while residues from the neighbouring protomer, including Pre9 K516 and F519, further contribute to PI binding (**Fig. 3b**). Hydrophobic interactions along Pre9 helix involving F519 and V520, together with disulfide bonds formed by C474 and C477 from TM8, can also be found at the dimer interface (**Fig. 3c, d**).

Notably, the CTL1 homodimer displays a slight asymmetry, with one side exhibiting closer packing between the TM3/TM4 and TM5/TM6 hairpins (**Fig. 3a**). Consistent with this, 3D variability analysis^38^ (3DVA) revealed continuous motions centred on TM3-TM6 and the TM8/Pre9 dimerisation interface (**Supplementary Fig. 9**). The gradual loss of density in these regions suggests intrinsic flexibility within the CTL1 dimer.

To investigate *in vivo* CTL1 dimerisation, we performed NanoBiT complementation assays using N- and C-terminally tagged CTL1 constructs, all of which were expressed at similar levels (**Supplementary Fig. 10a, b**). All complementary NanoBiT pairs generated robust luminescence above background, confirming CTL1 homodimerisation *in vivo* (**Fig. 3e**). Signal intensity varied depending on the orientation of the NanoBiT fragments (**Fig. 3e**), suggesting that steric factors or conformational constraints may influence either CTL1 dimer formation or the efficiency of NanoBiT fragment reconstitution. Importantly, C474S and C477S substitutions significantly reduced dimerisation, whereas the Pre9 double mutant F519A/V520A nearly abolished NanoBiT complementation without affecting protein expression (**Fig. 3f and Supplementary Fig. 10c**), suggesting a role in CTL1 dimerisation.

### CTL1 is a selective phospholipid-binding protein

A remarkable feature of CTL1 cryo-EM structure is the abundance of endogenous lipids bound at its transmembrane periphery and to a large amphipathic central cavity exposed to the outer leaflet (**Fig. 4a, d**). Although our analysis focused on the higher-resolution monomeric structure, equivalent lipid densities are observed at corresponding positions in the dimer. Notably, two non-protein densities that resemble phospholipid-like molecules were found within the exposed cavity in the cytosolic leaflet occupying two distinct binding sites referred to as position 1 and position 2 (**Fig. 4a**, **right**).

**Figure 4.**
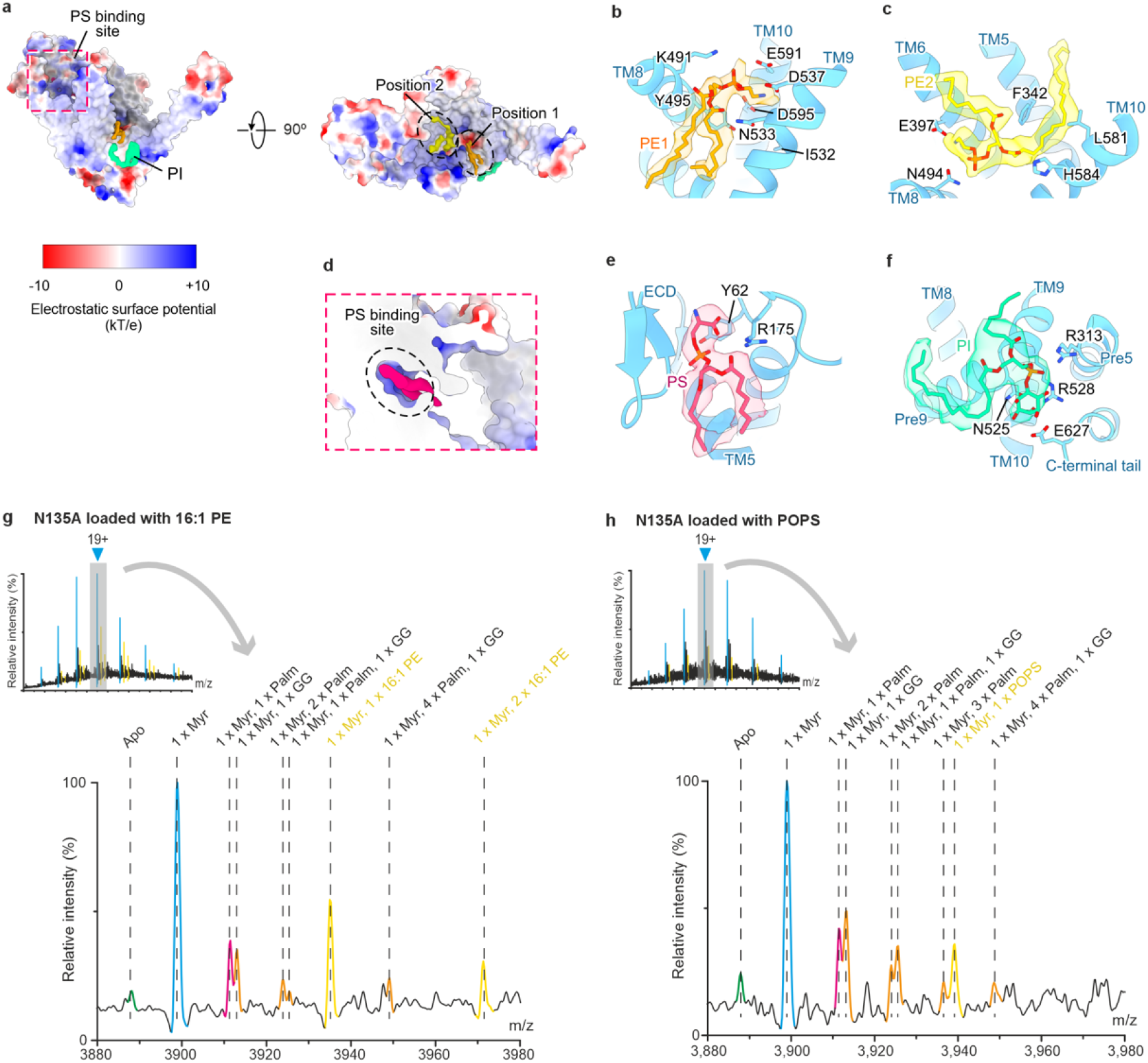
CTL1 interacts with phospholipid molecules. **a** Electrostatic surface potential of CTL1 with the PI density shown in light green, highlighting the PS-binding site (pink dashed line) in the front view. Right, top view showing the negatively charged binding positions 1 and 2 occupied by PE1 (orange) and PE2 (yellow) densities, respectively. **b-c** Close-up views of positions 1 (**b**) and 2 (**c**), showing PE1 (orange) and PE2 (yellow) molecules within their corresponding cryo-EM densities, with interacting residues shown as sticks. **d** Cross-sectional view of the PS-binding site outlined in **a** (pink dashed box), showing the positively charged binding pocket with the PS density depicted in deep pink. **e** Close-up view of the PS-binding pocket showing a PS molecule fitted into the corresponding density with interacting residues shown as sticks. **f** Close-up view of the PI density indicated in panel **a** (light green), showing a PI molecule within its density and interacting residues shown as sticks. **g, h** nMS of CTL1-N135A reveals binding of 16:1 PE (**g**) and POPS (**h**). Mass shifts corresponding to 16:1 PE (**g**) and POPS (**h**) are indicated. Lipid adduct peaks are highlighted yellow.

To further define the identity of these co-purified ligands with CTL1, we employed lipidomic analysis and nMS, which identified the binding of two molecules of 16:1 PE (**Fig. 4g**; **Supplementary Fig. 7d**). This observation, together with the shape of the densities and the electrostatic environment of the binding pockets, supported the assignment of both ligands as PE molecules (**Fig. 4b, c**). The lateral PE1 site is lined by conserved residues (**Supplementary Fig. 11 and Supplementary Table 1**) from TM8-TM10 (K491, N533, D537, E591 and D595) with Y495 and I532 lining the base of the binding pocket. In contrast, the central PE2 molecule interacts with residues from TM5 (F342), TM6 (E397), TM8 (N494), and TM10 (L581 and H584). In agreement with these structural observations, both total PE extracts and purified POPE significantly increased CTL1-GFP thermostability, further supporting an interaction between CTL1 and PE (**Supplementary Fig. 12a, b**). Notably, among the residues lining the PE1 site, D537 and D595 displayed the closest interactions with the ethanolamine headgroup (**Fig. 4b**). Consistent with this structural observation, alanine substitution of these two residues produced the most pronounced effects, with both D537A and D595A reducing POPE-mediated CTL1 thermostabilisation without significantly affecting PS and PI binding (**Supplementary Fig. 12a, c**). The agreement between the structural contacts and the functional effects of these substitutions supports a direct role for D537 and D595 in PE recognition and CTL1 stabilisation.

Intriguingly, lipidomics and nMS further detected binding of 1-palmitoyl-2-oleoyl-phosphatidylserine (POPS) (**Fig. 4h**). Inspection of the ECD revealed a well-defined phospholipid density bound within a positively charged pocket, consistent with PS, with its headgroup coordinated by Y62 and R175 (**Fig. 4d, e**). Unlike PE, which occupies the central cavity, PS binds at the extracellular boundary. In agreement, POPS also significantly stabilised CTL1-GFP during thermal denaturation, supporting the notion that PS binding contributes to CTL1 structural stability (**Supplementary Fig. 12a, b**). Notably, Y62 and R175, which directly coordinate the PS headgroup in the structure, were also functionally required for this interaction. The Y62A/R175A variant substantially reduced POPS-mediated thermal stabilisation compared with wild-type CTL1, establishing a direct correspondence between the structural PS-binding site and its functional contribution to CTL1 stabilisation (**Supplementary Fig. 12a, c**).

In addition, consistent with the structural assignment of PI at the CTL1 dimer interface, a corresponding weaker PI density was also observed in the monomer, likely due to the absence of the adjacent protomer which appears to stabilise PI binding in the dimer (**Fig. 4f**). In agreement with these observations, 1-palmitoyl-2-oleoylphosphatidylinositol (POPI) also enhanced CTL1-GFP thermostability (**Supplementary Fig. 12a, b**). By contrast, and consistent with lipidomic analysis, both 1-palmitoyl-2-oleoylphosphatidylcholine (POPC) as well as total PC extracts failed to confer detectable thermal stabilisation (**Supplementary Fig. 12a, b**), suggesting that CTL1 selectively recognises specific phospholipid headgroups rather than interacting non-specifically with membrane phospholipids. In addition, total egg PA enhanced CTL1-GFP thermostability, consistent with the enrichment of PA species in lipidomic analysis and supporting our tentative assignment of surface-exposed densities on CTL1 as PA based on their shape (**Supplementary Fig. 8d and 12a, b**).

### CTL1 induces local bilayer deformation in molecular dynamics simulations

The conical architecture of CTL1 observed in the cryo-EM structure (**Supplementary Fig. 13**), together with the presence of phospholipid densities within its central cavity, suggested that the protein may locally deform the surrounding membrane to facilitate lipid recruitment into the protein core. Notably, the detergent micelle surrounding CTL1 narrows from the extracellular surface towards the central cavity, by approximately 17 Å in the monomer and 25 Å in the dimer, creating a funnel-like constriction immediately above the two PE-binding sites (**Supplementary Fig. 13a**). Although this detergent-micelle geometry does not by itself demonstrate membrane deformation, it provided a structural rationale to test whether CTL1 can affect lipid bilayers.

To address this, we performed all atomistic MD simulations of monomeric and dimeric CTL1 embedded in a POPC:POPE:POPS:CHOL (5:2:1:2) symmetric membrane. This composition was selected as a defined vesicle-mimetic membrane containing major phospholipid classes relevant to membrane organisation, vesicle dynamics, and membrane fusion^39^. PC/PE/PS-containing membranes supplemented with cholesterol have been used to investigate the physical properties of vesicle membranes and their propensity to undergo membrane fusion, providing a tractable model for studying membrane reshaping processes^39^. The symmetric composition was chosen to isolate the effects of CTL1 on membrane deformation and phospholipid recruitment, without introducing additional variables associated with leaflet-specific lipid distributions.

Whereas the protein-free membrane remained essentially flat and unaltered, CTL1 induced pronounced local membrane deformation, particularly in the dimer (**Fig. 5a**), consistent with what we observed in the cryo-EM maps of CTL1 in micelles. Specifically, the monomer produced an average local membrane compression of -4.04 Å, whereas the dimer generated a much broader region of hydrophobic mismatch, resulting in an average membrane compression of -5.41Å (**Fig. 5b**), that propagated into the surrounding bilayer and imposed a characteristic saddle-shaped (negative Gaussian curvature) deformation (**Fig. 5a, c and Supplementary Fig. 13b**). The resulting deformation brings the outer leaflet into close proximity with the central cavity, providing a structural mechanism by which membrane deformation could facilitate phospholipid recruitment into the protein core.

**Figure 5.**
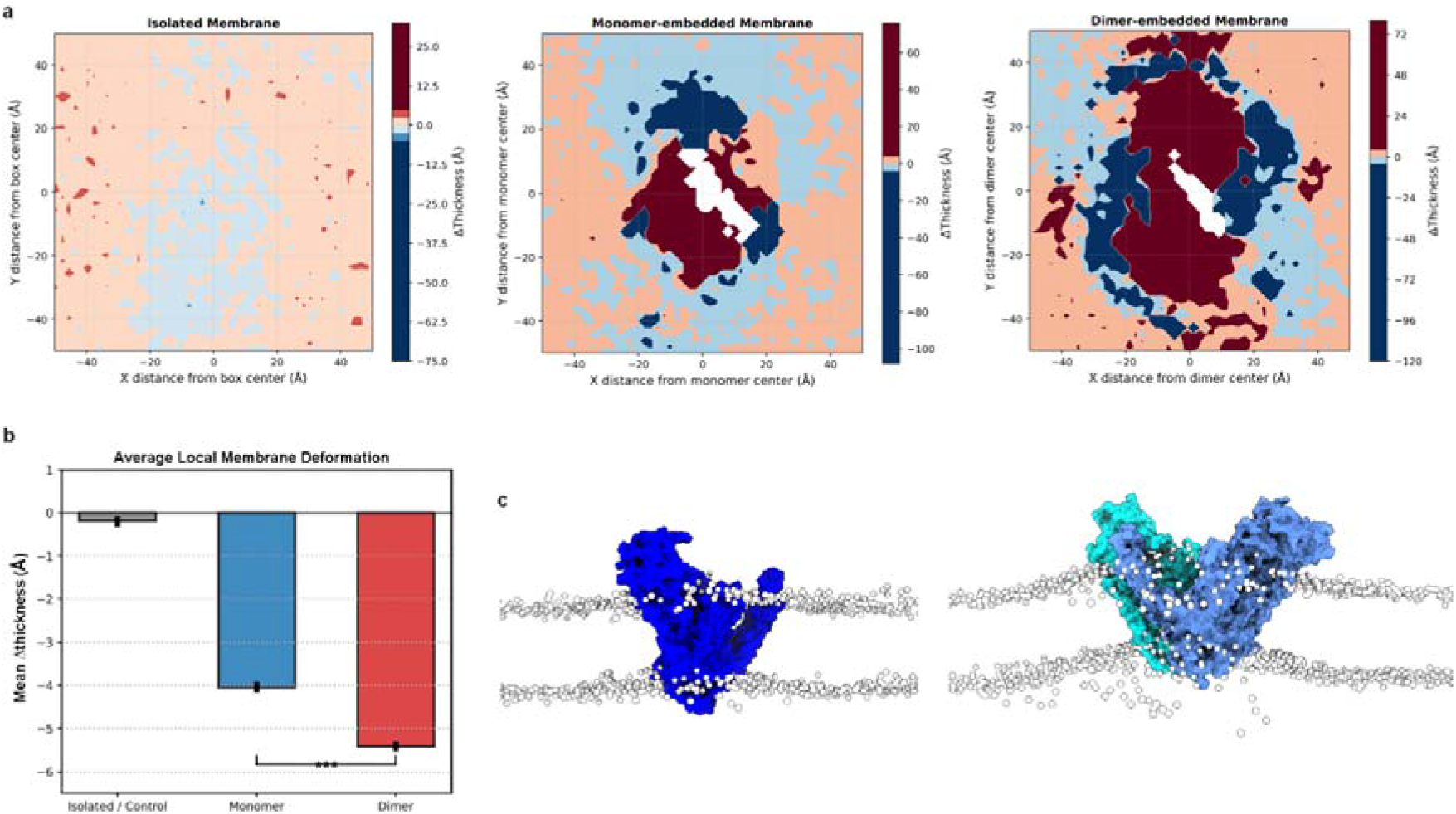
Membrane thickness changes induced by monomeric and dimeric CTL1. **a** Local bilayer thickness was calculated as the time-averaged distance between the phosphate headgroups of the upper and lower leaflets Z_upper_ - Z_lower_ on a 2.0 Å XY grid centred around the protein centre of mass (or box centre for the control). Thickness changes (ΔThickness) was calculated relative to the average bulk thickness of the isolated membrane (43.78 Å), with ΔThickness = 0 Å corresponding to the unperturbed membrane and negative values indicating membrane compression/thinning. Central masked patches correspond to protein core areas where no lipid phosphate headgroups are present. **b** Mean local thickness changes (ΔThickness) for the isolated control membrane (grey, -0.19 Å), monomer-embedded membrane (blue, -4.04 Å), and dimer-embedded membrane (red, -5.41Å). Negative values reflect a net average compression and thinning of the hydrophobic core. Error bars indicate the standard error of the mean (SEM). The asterisks (***) denote a highly statistically significant difference (*p* < 0.001) between the monomeric and dimeric systems. A Two-Sample Welch’s *t*-test was used to o determine the statistical significance of the distinct deformation gap between the monomeric and dimeric states. **c** Representative snapshots from MD simulation showing surface representations of monomeric (left) and dimeric (right) CTL1 embedded in a phospholipid bilayer (phosphorus atoms shown as white spheres). Whereas the monomer induced only limited deformation (left), the dimer induced symmetric membrane depressions along its horizontal axis and corresponding elevations along the perpendicular axis (right), wrapping the host bilayer into a distinct saddle-shaped geometry (Negative Gaussian Curvature).

### Molecular determinants of phospholipid recruitment

CTL1 structural analysis identified a cluster of highly conserved polar and positively charged residues (**Supplementary Fig. 11 and Supplementary Table 1**) extending from the extracellular vestibule toward the PE1-binding site (K454, R457, R476, K480, K491, K544, H584) (**Supplementary Fig. 14a)**. The presence of this polar network within the membrane interior is unusual and suggests that it may contribute to phospholipid coordination rather than serve a structural function. To investigate this possibility, we further analysed the MD simulations of dimeric CTL1 embedded in the model membrane. Throughout the simulations, phospholipid molecules from the extracellular leaflet entered the central cleft and migrated toward positions resembling those observed in the cryo-EM structure (**Fig. 6a**; **Supplementary Movie 1**).

**Figure 6.**
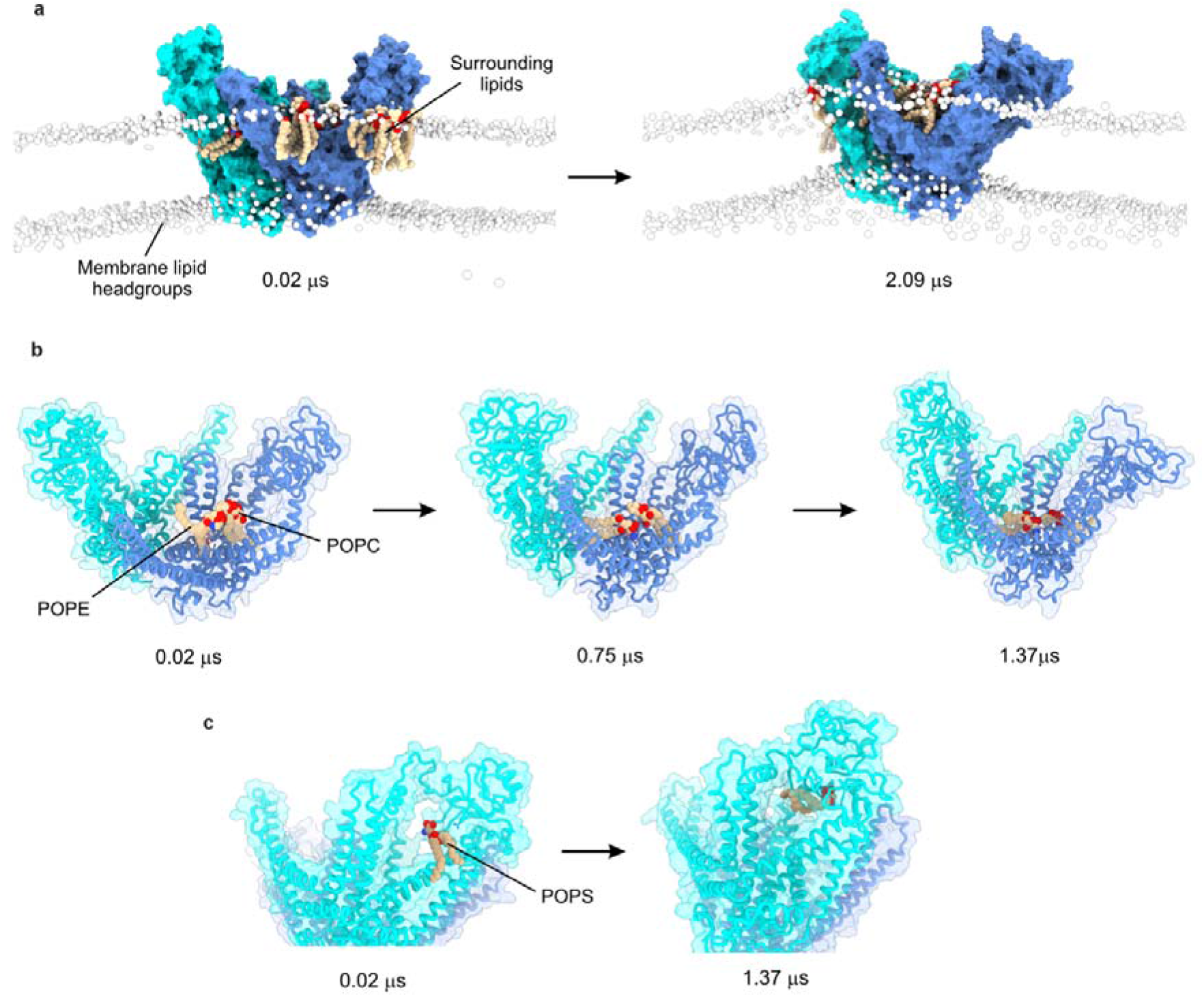
Lipid recruitment and binding to dimeric CTL1. **a** Snapshots from MD simulations showing dimeric CTL1 recruiting surrounding phospholipids from the bulk membrane lipids (white spheres). Surface representations of the two protomers are shown in cyan and blue. **b** Snapshots from MD simulations in which two phospholipid molecules are recruited sequentially into binding positions 1 and 2 within the central cavity of CTL1, shown as a cartoon representation. **c** Snapshots showing a POPS molecule entering the PS-binding pocket located in the ECD during the simulation.

Notably, a POPE molecule rapidly occupied the PE1 site and remained stably bound throughout the simulations, reproducing the cryo-EM binding mode (**Fig. 6b**; **Supplementary Movie 2**). In contrast, lipids within the PE2 site remained substantially more dynamic, sampling multiple orientations, suggesting that PE2 represents a lower-affinity recruitment intermediate compared with the more selective PE1 pocket. Likewise, although POPS constituted only a minor fraction of the model membrane, a POPS molecule specifically entered the positively charged pocket within the ECD early in the simulations and remained stably coordinated by Y62 and R175 throughout consistent with the cryo-EM structures and thermostability assays, supporting the existence of an extracellular PS-recognition site (**Fig. 6c**; **Supplementary Fig. 12a, c; Supplementary Movie 3**).

Remarkably, lipid recruitment was accompanied by local rearrangements within the TM core that promoted hydration of the transmembrane region, generating transient water-accessible pathways extending toward the central cavity. Water penetration was associated with subtle conformational rearrangements involving the TM2, TM9, and TM10 region allowing water molecules to interact with residues of the conserved polar network (**Supplementary Fig. 11b and 14b)**, regulated by two hydrophobic residues, W487 and F541, which act as dynamic gates controlling solvent access. Consistent with a functional role of this network in PE coordination, the CTL1-GFP sextuple variant targeting the polar network (R457A/W487A/E490A/K491A/F541A/K544A), which displayed thermal stability comparable to that of wild-type CTL1, showed a marked reduction in POPE-mediated stabilisation (**Supplementary Fig. 12a, c**). Notably, this effect was selective for PE, as the sextuple mutant retained POPS- and POPI-mediated thermostabilisation. Thus, disruption of the conserved polar network does not cause a general impairment of CTL1-lipid interactions but specifically compromises the ability of PE to stabilise the protein. Together, these findings support a collective and lipid-selective contribution of the conserved polar network to PE coordination and its progression through the protein cavity.

Overall, the MD simulations support a model in which CTL1 recruits phospholipids from the outer membrane leaflet into a conserved central cavity while transiently hydrating its TM core. Although the precise molecular mechanism remains unresolved, all these findings provide a mechanistic framework for CTL1-mediated phospholipid recognition and membrane deforming processes.

## Discussion

More than two decades after its identification^6^, the molecular basis of CTL1/SLC44A1 function remains elusive. Although CTL1 was initially proposed to function as a Na^+^-independent choline transporter, this long-standing model has been increasingly challenged by the recent identification of FLVCR1^40^ and FLVCR2^34^ as the primary Na^+^-independent choline transporters, together with inconsistent evidence that CTL1 homologues transport choline^25,41^. Here, our thorough functional analysis confirmed negligible CTL1-mediated choline uptake (**Fig. 1 and Supplementary Fig. 1**). By combining cryo-EM, nMS and lipidomics, we identify endogenous phospholipid ligands of CTL1 and uncover a structurally defined phospholipid-recognition environment within the protein, characterised by conserved polar and charged residues that coordinate phospholipid headgroups.

Multiple phospholipid-binding sites were identified throughout the cryo-EM structure, including PE within the transmembrane cavity, PS within the ECD, PI at the dimer interface and PA at the membrane-facing surface, with nMS, lipidomic and thermostability analysis independently supporting preferential interactions with these lipid species. Numerous contacts with endogenous cholesterol and a putative P-LPC further highlight the diversity of lipids associated with CTL1. The presence of the latter is particularly notable given the abundance of plasmalogens in myelin, where CTL1 is enriched and its dysfunction has been associated with defects in myelination and neurological function^16,17,20^. The association of CTL1 with PS at the ECD is also noteworthy given that PS is normally restricted to the cytoplasmic leaflet^42,43^. Its exposure at the cell surface is typically linked to apoptosis, platelet activation and inflammatory signalling, raising the possibility that PS recognition by CTL1 may have a functional significance beyond structural lipid binding that needs further investigation.

MD simulations further suggest that CTL1 combines selective phospholipid binding with local membrane deformation. In our simulations, CTL1 induces local membrane thinning and curvature, creating a funnel-shape deformation that may facilitate phospholipid recruitment into its central cavity, with this effect enhanced upon oligomerisation (**Fig. 5**, **Supplementary Fig. 13**). The distinct dynamics at PE1 and PE2 sites may also suggest a possible differential or sequential route for phospholipid entry into the experimentally observed binding sites (**Fig. 6a**; **Supplementary Movie 1**). The membrane deformation observed in these simulations is consistent with the conical architecture of CTL1, particularly in its dimeric state. A relevant precedent is provided by the SARS-CoV-2 M protein, in which specific lipid interaction stabilises a cone-shape M conformation, promoting local membrane bending and thinning during viral assembly and budding^44,45^. Membrane reshaping is accompanied by enrichment of particular phospholipids around the transmembrane region of M^46^, suggesting an association between membrane deformation and local lipid organisation. In CTL1, the combination of a conical architecture, phospholipid binding and membrane deformation observed in the simulations raises the possibility that a similar physical mechanism may operate. The conical geometry of CTL1 could favour local membrane bending and enrichment of particular phospholipids around the transmembrane region, potentially facilitating their entry into the transmembrane cavity and occupancy of the phospholipid-binding sites observed by cryo-EM. This model would provide a potential link between the lipid interactions identified here and the membrane-reshaping behaviour observed in MD.

CTL1 is also subject to several lipid post-translational modifications, including geranylgeranylation, myristoylation and palmitoylation. Lipidation can influence membrane targeting, trafficking and protein organisation by providing hydrophobic membrane contacts or altering the interaction of proteins with specific lipid environments^47^. Prenylation and myristoylation, for example, can regulate membrane association and intracellular trafficking, whereas palmitoylation can additionally control membrane-domain partitioning and the organisation of membrane proteins within cholesterol- and sphingolipid-enriched domains^47,48^. Consistent with this possibility, CTL1 has been identified in lipid-raft proteomic datasets^49,50^, and our cryo-EM structures reveal extensive contacts with endogenous cholesterol around the transmembrane region. Palmitoylation can also influence how membrane proteins respond to the physical properties of the bilayer. For example, palmitoylated cysteine clusters in TMEM63B are required for membrane association and plasma membrane localisation and contribute to activation of lipid scrambling in response to changes in membrane curvature and thickness^51^. Together, these observations raise the possibility that lipid modifications contribute to the partitioning of CTL1 within specific membrane environments and modulate its interactions with the surrounding bilayer. Whether these modifications influence CTL1 trafficking, membrane-domain organisation or its response to local membrane properties remains to be determined.

By linking CTL1 to phospholipid recognition and membrane deformation, our model may also provide a framework for understanding the broader cellular consequences of CTL1 loss. CTL1 deletion has been associated with altered membrane lipid homeostasis^13,14,16,17,20^ and defects in intracellular trafficking^12^, including impaired delivery of membrane proteins to the plasma membrane in *Arabidopsis thaliana*^13,14^. Although these phenotypes have previously been interpreted primarily in the context of altered choline or phospholipid metabolism, they are also compatible with a role for CTL1 in maintaining membrane organisation. Membrane curvature, bilayer thickness and lipid packing can influence membrane-associated proteins, lipid-transfer processes and trafficking pathways, providing a potential link between CTL1-dependent lipid interactions and these cellular phenotypes. The recurrence of membrane-related phenotypes following CTL1 disruption across yeast^12^, plants^13,14^, mice^16,17^ and humans^20^ further raises the possibility that membrane reshaping and phospholipid interactions may represent a conserved aspect of CTL1 biology.

In summary, our findings support a view of CTL1 as a phospholipid-interacting membrane protein and we posit that its function is closely linked to lipid recognition and the physical properties of the surrounding bilayer. Structural, biochemical and computational analyses reveal a previously unrecognised capacity for phospholipid recognition and identify a potential role for CTL1 in shaping the local lipid environment and membrane organisation. More broadly, these findings raise the possibility that SLC44 proteins represent a distinct group of lipid-interacting membrane proteins and provide a molecular framework for understanding how CTL1-dependent membrane organisation contributes to physiological processes and to diseases associated with CTL1 dysfunction, including neurodegeneration, myelination defects and cancer-associated membrane adaptation.

## Methods

### Construct generation

Full length human CTL1 (*SLC44A1*, isoform 1) and FLVCR2 (*SLC49A2*, isoform 1) coding sequences were codon-optimised, *de novo* synthesised and cloned into the mammalian expression vector pcDNA3.1(+) (GenScript). The CTL1 sextuple (R457A/W487A/E490A/K491A/F541A/K544A) variant was generated by GenScript, upon request. The remaining CTL1 variants were generated using the primers listed in Supplementary Table 2. For protein expression and fluorescence microscopy assays, a C-terminal HRV3C cleavage site followed by a superfolded GFP, a Twin-Strep and a 10xHis tags was added in frame.

The LgBiT and SmBiT fragments of NanoLuc Luciferase (Promega) were amplified by PCR with glycine-serine linkers and cloned into N- or C-terminal gateway-compatible entry vectors. Using gateway technology, LgBiT and SmBiT fragments were added to either the N- or C-terminal domains of CTL1 in a pcDNA3.1(+) backbone. The integrity of all cloned constructs was confirmed by DNA sequencing.

### Expression and purification of CTL1

Recombinant CTL1 protein was expressed in HEK293F cells transfected with plasmid and FectoPro in a ratio of 1:1. Transfected cells were maintained in Freestyle 293 Expression medium (Thermo Fisher Scientific) at 37°C, 8% CO_2_ and 80% humidity in an incubator shaker. After 48 h of protein expression, cells pellet was collected and stored at -80°C. Cell pellet was resuspended with Dounce homogeniser in solubilisation buffer containing 50 mM Tris-HCl pH 7.4, 150 mM NaCl, 2 mM ATP/MgCl_2_, 0.5 mM PMSF, 1 mM benzamidine, DNAse I, 1% (w/v) n-Dodecyl-β-D-Maltoside (DDM) and 0.01% (w/v) cholesterol hemisuccinate (CHS) (Anatrace). The lysate was incubated for 2 h at 4°C with continuous rotation followed by high-speed ultracentrifugation at 300,000 x*g* for 30 min at 4°C. The resulting supernatant was loaded onto Strep-Tactin XT affinity column, then washed with 40 CV of buffer containing 50 mM Tris-HCl pH 7.4, 150 mM NaCl, 1 mM ATP/MgCl_2_, 0.04% DDM and 0.004% CHS, and finished with 10 CV of the same washing buffer without ATP/MgCl_2_. The protein was eluted by on-column HRV3C protease digestion, concentrated and further purified by size-exclusion chromatography (SEC) on a Superdex200 increase 10/300 GL column (Cytiva) with SEC buffer containing 50 mM Tris-HCl pH 7.4, 150 mM NaCl, 0.02% DDM and 0.002% CHS. CTL1 dimer and monomer peak fractions were pooled and concentrated with a 100 kDa MWCO Amicon to 3 mg/ml of CTL1 for cryo-EM sample preparation. The CTL1-N135A variant, designed to remove N135-glycosylation for nMS analysis, was expressed and purified in n-Nonyl-β-D-Glucopyranoside (NG) (Anatrace) following the same protocol.

### Cryo-EM sample preparation and data collection

Cryo-EM sample preparation was conducted using a Vitrobot Mark IV (ThermoFisher) at 4 °C and 100% humidity. 3 μL of the purified protein was applied onto a glow-discharged UltrAuFoil® R 0.6/1 300 mesh grid (Quantifoil), blotted for 4s and plunge-frozen in liquid ethane. Cryo-EM data were acquired on a 300kV Titan Krios G4 (Thermo Scientific), equipped with a BioContinuum energy filter and a K3 direct electron camera (Gatan) operating in super resolution mode at x130.000 nominal magnification and a calibrated pixel size of 0.6462 Å/pix. Movies were automatically collected using EPU2 (Thermo Scientific), with Aberration Free Image Shift (AFIS) and Fringe-free imaging (FFI), at a defocus range of -1.0 μm to -1.8 μm with a total exposure of 60.7 e^-^/Å^2^ dose-fractioned into 61 frames per stack.

### Cryo-EM data processing

Two independent exposure datasets were collected, comprising 31,562 and 25,687 micrographs, respectively. Initial frame alignment and contrast transfer function (CTF) estimation were performed using cryoSPARC Live. Micrographs with an estimated CTF resolution better than 4 Å were accepted for further processing. Initial particle picking was conducted using blob picking mode in cryoSPARC Live followed by 2D classification. Initial 2D classes, showing features of the extracellular region and transmembrane domains within a discoidal micelle, were selected as templates for subsequent automatic particle picking. A total of ∼5.7 million particles were picked from each batch and subjected to multiple rounds of 2D classification. Notably, some 2D class averages that resemble dimeric CTL1 were clearly visible, displaying larger micelles with symmetrical features in both extracellular and transmembrane region.

Particles from selected 2D class averages were used to generate *ab initio* 3D reconstructions with five classes, starting at a maximum resolution of 8 Å and gradually increased to 4 Å to obtain higher resolution. *Ab initio* models of both CTL1 monomer and dimer were successfully obtained, confirming a substantial proportion of dimeric CTL1 in the sample. These *ab initio* models serve as initial references for subsequent rounds of heterogenous refinement. Discarded 2D classes were used to generate decoy models to trap poor quality particles and clean the dataset.

In parallel, particles from previously selected 2D classes were used as templates for Topaz^52^ training. Automatic Topaz picked particles were subjected to multiple rounds of heterogenous refinement with initial resolution of 8 Å, using previously obtained *ab initio* models as initial references. The resulting 3D models of CTL1 monomer and dimer from both datasets reached nominal resolutions of 5-6 Å. To improve resolution, we next merged both datasets and then divided in monomer and dimer particles subsets, yielding 1,813,776 monomeric and 1,180,561 dimeric particles. These particles subsets independently were subjected for new rounds of heterogenous classification. The best 3D class of CTL1 monomer, containing 515,168 particles, was subjected to non-uniform refinement with an initial low pass filter of 15 Å, obtaining a final map at 3.05 Å nominal resolution according to the gold-standard Fourier shell correlation (FSC). Similarly, the dimeric CTL1 reconstruction, comprising 373,713 particles, was non-uniform refined by applying C2 symmetry with relaxed symmetry maximisation mode, yielding a final cryo-EM map at 3.23 Å nominal resolution according to FSC.

To analyse the conformational heterogeneity of dimeric CTL1, 3D variability analysis^38^ (3DVA) was performed using 242,000 particles and a mask was applied to the protein to exclude the detergent micelle. The analysis was run using three variability components and a filter resolution of 6 Å. The first component was subjected to a 3DVA display job to visualise continuous motions.

### Model building of CTL1

An initial model of the CTL1 monomer was generated using ModelAngelo^53^ and then fitted into the monomeric 3.05 Å resolution cryo-EM map in UCSF ChimeraX^54^. The dimeric CTL1 model was subsequently generated using UCSF ChimeraX by rigid-body fitting of two monomeric units into the 3.23 Å dimeric map. The first 15 N-terminal residues and the last 29 C-terminal residues could not be modelled due to the absence of defined density. The protein Cα backbone and side chains were manually inspected and adjusted in Coot^55^. Additional features, including N-glycan, geranylgeranylation on cysteines, Mg^2+^ ion, cholesterol and phospholipid molecules were also manually added and refined in Coot. The identity of the Mg^2+^ ion was validated using the server CheckMyMetal (https://cmm.minorlab.org)^56^. Coordinates and geometry restraints for plasmalogen lysophosphatidylcholine (P-LPC) were generated using eLBOW in Phenix^57^. The resulting monomeric and dimeric CTL1 models were automatically refined against their respective cryo-EM maps using real-space refinement in Phenix^58^ to optimise model geometry. All the figures were prepared with UCSF ChimeraX.

### Cell culture and transfection

HeLa, HEK293T, HCT116, CHO and MDCK cells were maintained at 37°C in a humidified 5% CO_2_ environment in Dulbecco’s modified Eagle’s medium supplemented with 10% fetal bovine serum, penicillin (50 U/ml), streptomycin (50 μg/ml), and 2 mM L-glutamine. For uptake assays, all cell lines were transiently transfected using Lipofectamine 2000 (Invitrogen), while HEK293T cells used for dimerisation assays were transfected using Transfectin Lipid Reagent (Bio-Rad) following the manufacturer’s protocol. Experiments were conducted 24 to 48 h post-transfection. All cell lines tested negative for *Mycoplasma* contamination.

### Western blot analysis

Cells were lysed in lysis buffer containing Tris-HCl 20 mM pH 7.4, NaCl 150 mM, Triton X 100 0.1%, sodium orthovanadate 0.1 mM, PMSF 0.5 mM, supplemented with protease inhibitors (Roche). After centrifugation at 21,000 x*g* for 10 min, the supernatants were mixed with Laemmli sample buffer, heated at 56 °C for 5 min and subjected to SDS-PAGE and Western blot analysis. Anti-CTL1 antibody from Proteintech (14687-1-AP) with 1:1,000 dilution was used. Images were acquired using ChemiDoc MP system and its accompanied software Image Lab Touch.

### Split-nanoluciferase experiments

NanoBiT complementation assay was performed using constructs in which CTL1 was fused to either the Small BiT (sb) or Large BiT (lb) luciferase fragments. Both fragments were introduced at either the N- or C-terminus of CTL1, generating four distinct constructs, which showed similar expression levels: sb-CTL1, CTL1-sb, lb-CTL1, and CTL1-lb. Forty-eight hours after transfection of HEK293T with a 1:1 ratio of SmBiT- and LgBiT-tagged CTL1 constructs, cells were detached, washed with PBS and resuspended into 1 ml of PBS. 50 μl of the suspension was seeded into white, flat-bottom 96-well plates (Bio-Rad; 50,000 cells/cm^2^), and NanoLuc and Rluc activities were measured using Promega reagents according to the manufacturer’s instructions. The transfection control of pGK-Rluc was included in all conditions. A Large BiT-containing vector was used as a control to check non-specific SmBiT-LgBiT interactions. Luminescence was recorded with a CLARIOstar plate reader after 10 min of incubation, and the normalised NanoLuc signal (normalised light unit/relative light unit) was used for analysis. Luciferase fold increase was calculated by dividing the luminescence collected for the tested combinations by the luminescence obtained for the corresponding negative control. Data are expressed as the mean ± SD of at least three experiments performed on different days and batches of cells.

### Choline transport assay in cells

Choline uptake was measured in pcDNA3.1-CTL1, pcDNA3.1-FLVCR2, and mock-transfected HeLa, HEK293T, HCT116, CHO and MDCK cells, 24 h post-transfection, as previously described^36^, by exposing replicate cultures at room temperature to [^3^H]-labelled choline chloride (1 μCi/ml; PerkinElmer) and unlabelled choline chloride added to the desired final concentration in transport buffer [10 mM HEPES (pH 7.5), 137 mM NaCl, 5 mM KCl, 2 mM CaCl_2_, and 1 mM MgSO_4_]. Initial rates of transport were determined using an incubation period of 1 min. Assays were terminated by washing with an excess volume of ice-cold stop buffer (5 mM choline chloride in transport buffer). To assess Na^+^ dependence, the buffer was supplemented with either 137 mM NaCl (Na^+^-containing conditions) or 137 mM NMDG (Na^+^-free conditions). To evaluate pH dependence, 10 mM HEPES was substituted by 10 mM MES. The pH of Na^+^-free buffers was adjusted with 10 N KOH, whereas NMDG-containing buffers were adjusted with 5 N HCl. Data are expressed as the mean ± SD of three experiments performed on different days and batches of cells.

### Visualisation of CTL1 by fluorescence microscopy

To analyse CTL1-GFP protein expression and plasma membrane localisation, fluorescence microscopy was performed on a semiconfluent monolayer of transfected cells cultured on glass coverslips, 24 h post-transfection. Glass coverslip-grown cells were rinsed three times with Hank’s Balanced Salt Solution (HBSS) containing Ca^2+^ and Mg^2+^ and incubated with 5μg/ml wheat germ agglutinin (WGA) labelled with Texas-Red (Thermo Fisher Scientific) at 37°C for 10 min, rinsed three times with HBSS-Ca^2+^-Mg^2+^ and fixed for 15 min in 4% paraformaldehyde. Fixed cells were then rinsed three times with phosphate-buffered saline and then mounted with aqua-poly/mount coverslipping medium (Polysciences Inc.). Images were taken using a Zeiss LSM980 laser scanning confocal microscope. All images were captured during 200 ms.

### CTL1 reconstitution into total brain lipids liposomes

Total porcine brain lipids (Avanti Polar Lipids Inc) were dried under N_2_ for at least 3 h to form a lipid film. Lipids were re-hydrated for 30 min at room temperature (RT) in reconstitution buffer containing 20 mM Tris-HCl (pH 7.4), 150 mM KCl and 1 mM CaCl_2_, and further bath-sonicated for 5 min. Ten freeze-thaw cycles were performed before extrusion 21 times through polycarbonate 400 nm filters to give unilamellar liposomes.

For protein incorporation, incremental additions of 10% Triton X-100 (w/v) aliquots were performed and monitoring of absorbance at 540 nm was used to detect liposome destabilisation. Then, DDM-purified CTL1 was added to a 1:40 protein/lipid ratio (w/w) and gently rotated at room temperature for 30 min before detergent removal by the addition of 40 mg/ml SM2 Bio-Beads per ml of sample. Bio-Bead additions were performed as follows: 2 h at RT, 1 h at 4°C, overnight at 4 °C, 2 h at RT and 2 h at RT. Bio-Beads were removed by gravity filtration and the sample was used immediately for assay measurements. For all reconstitution procedures, empty liposomes lacking any purified protein were generated using the same protocols.

To check liposome reconstitution efficiency, an aliquot of the resulting proteoliposome mixture was mixed with 80% sucrose (w/w) in reconstitution buffer to render a 50% sucrose solution. Sample was then overlaid with 2 ml of 40%, 30%, 20%, 10% and 5% sucrose and reconstitution buffer, and centrifuged at 185,000 x*g* for 19 h at 4°C with minimal acceleration and deceleration. Fractions (1 ml) were collected and analysed by SDS-PAGE.

### Choline transport assays in proteoliposomes

Choline uptake assays in proteoliposomes were performed as previously described^36,59^. In brief, choline uptake was initiated by mixing 20 μl of ice-cold (proteo)liposomes with 180 μl of transport buffer containing 50 μM [^3^H]-choline chloride (2 μCi/ml; PerkinElmer) in 10 mM HEPES (pH 7.4), 150 mM NaCl, 2 mM CaCl_2_, and 1 mM MgSO_4_.

Transport assays were carried out at room temperature for the indicated time periods and terminated by the addition of 2 ml of ice-cold stop buffer (5 mM choline chloride in transport buffer), followed by rapid filtration through 0.45-μm pore-size membrane filters (Sartorius Stedim Biotech). Filters were washed twice with 2 ml of cold stop buffer, dried, and the retained radioactivity was quantified by scintillation counting. Data are presented as mean ± SD from triplicate measurements at each time point in one representative experiment. A second independent experiment, performed on a different day using separate liposome reconstitutions, gave similar results.

### Thermal shift assay

CTL1 thermal shift assay was performed as previously described^59^ in DDM-solubilised and NG-delipidated wild-type or variant CTL1-GFP proteins^60^. GFP-fused CTL1 proteins were solubilised in 50 mM Tris-HCl pH 7.4, 150 mM NaCl, 0.5 mM PMSF, DNAse I, and 1% (w/v) DDM ON at 4°C with continuous rotation followed by high-speed ultracentrifugation at 300,000 x*g* for 30 min at 4°C. The clarified supernatant was loaded onto a Strep-Tactin XT affinity column, washed with 60 CV of buffer containing 50 mM Tris-HCl (pH 7.4), 150 mM NaCl and 0.4% (w/v) NG. The eluate was concentrated using a 100-kDa molecular weight cut-off Amicon centrifugal filter and further purified by size-exclusion chromatography on a Superdex 200 Increase 5/150 GL column (Cytiva) equilibrated with 50 mM Tris-HCl (pH 7.4), 150 mM NaCl and 0.4% (w/v) NG.

For thermal stability measurements, 150 µl of 0.03 mg/ml NG-purified wild-type or mutant CTL1-GFP was incubated for 10 min at temperatures ranging from 4 to 75 °C. Samples were subsequently centrifuged at 21,000 x*g* for 30 min at 4 °C to remove aggregated protein. An aliquot (120 µl) of the clarified supernatant was transferred to a flat-bottom 96-well plate, and GFP fluorescence was measured using a CLARIOstar microplate reader (BMG Labtech). Melting temperatures (*T_m_*) were determined by fitting the fluorescence signal, normalised to the fluorescence measured at 4 °C, to a sigmoidal dose-response curve using GraphPad Prism (GraphPad Software).

To evaluate the effect of ligands on CTL1 stability, 150 µl of 0.03 mg/ml NG-purified wild-type or mutant CTL1-GFP were mixed with an equal volume of NG-solubilised phospholipids (5 mg/ml) or the corresponding vehicle control and incubated for 60 min at 4°C. Phospholipids were prepared as previously described^60^. Samples were then heated for 10 min at a temperature 5 °C above the corresponding *T_m_* of each CTL1 construct, followed by centrifugation at 21,000 x*g* for 30 min at 4 °C. An aliquot (120 µl) of the clarified supernatant was transferred to a flat-bottom 96-well plate, and GFP fluorescence was measured using a CLARIOstar microplate reader. Ligand-induced stabilisation was expressed as the fluorescence remaining after heat treatment relative to the corresponding vehicle control. Data are presented as the mean ± SD of three independent experiments performed on different days using independently purified CTL1 protein preparations.

### Sequence alignment

Evolutionary conservation across CTL1 homologs was assessed using ConSurf^61^, and the 15 homologs with the highest sequence similarity were selected for multiple sequence alignment in PROMALS3D^62^, based on both amino acid sequence and secondary structure information. Sequence alignment of human CTL family members (CTL1-5) was performed in Jalview^63^, which was also used for visualisation of all sequence alignments and figure preparation.

### Native mass spectrometry analysis

Purified CTL1 (wild-type and N135A) was buffer-exchanged into 500 mM ammonium acetate, pH 7.5, containing 0.4% (w/v) NG using a Zeba Spin 40K MWCO desalting column (Thermo Scientific). Protein concentration was determined using a NanoDrop One and adjusted to the desired concentration for native mass spectrometry analysis. Typically, 3 μL of desalted protein was loaded into an in-house prepared gold-coated capillary and electrosprayed into a Q Exactive UHMR hybrid quadrupole-Orbitrap mass spectrometer (Thermo Fisher Scientific) using the following settings: capillary voltage, 1.4 kV; UHV pressure, 7.5 × 10^−10^ mbar; capillary temperature, 200 °C; and resolution, 17,500. Unless otherwise stated, all experiments were performed in positive mode. Raw MS data were processed with Xcalibur 4.1.50 (Thermo Fisher Scientific), and spectra were deconvoluted using UniDec v6.0.1^64^.

Hydroxylamine treatment was used to reduce cysteine modification on wild-type CTL1 using a modified protocol^65^. Purified CTL1 was incubated with 0.5 M hydroxylamine overnight at 4°C and subsequently buffer-exchanged into buffer containing 500 mM ammonium acetate (pH 7.5) and 0.4% NG. nMS was collected similarly.

For lipid-loading experiments, the glycosylation-deficient N135A CTL1 mutant was used to improve spectral quality and facilitate the interpretation of lipid-bound species. POPS, POPE and 16:1 PE (Avanti) were dissolved in 500 mM ammonium acetate buffer containing 0.4% DM (5xCMC) at a concentration of 0.2 mM. Typically, 44 μM CTL1 was incubated with 100 μM lipids on ice for 10 min before injection into the mass spectrometer. Raw data were analysed similarly as above.

### Lipidomics

Lipids were extracted from purified CTL1 samples or from flowthrough fractions using a previously described methyl-tert-butyl ether (MTBE)/methanol extraction method^66^. The extracted lipids were dried and re-suspended in buffer (80% lipidomics buffer A/20% lipidomics buffer B). For LC-MS/MS analysis, lipids were separated on a C18 column (Acclaim PepMap 100; 75 µm × 150 mm, 3 µm particle size; Thermo Fisher Scientific) using a Dionex Ultimate 3000 UHPLC system. A binary solvent system was used: buffer A (60% acetonitrile, 40% water, 10 mM ammonium formate, and 0.1% formic acid), buffer B (90% isopropanol, 10% acetonitrile, 10 mM ammonium formate, and 0.1% formic acid). Eluted lipids were analysed on an Orbitrap Eclipse Tribrid mass spectrometer (Thermo Fisher Scientific) operated in both positive and negative mode using data-dependent acquisition. Lipidomics data were processed and analysed in MS-DIAL-5.4^67^.

### Proteomics

An SDS-PAGE gel band for purified CTL1 was sliced, destained, reduced with 50 mM TCEP for 10 min at 60 °C, alkylated with 100 mM iodoacetamide (IAA) at room temperature for 30 min in the dark, and incubated with trypsin at 37 °C overnight for proteolytic digestion. Digested peptides were dried using a SpeedVac vacuum concentrator (Thermo Fisher Scientific) and re-solubilised with 100% H_2_O with 1% formic acid for LC-MS/MS analysis. The tryptic peptides were analysed on a Dionex Ultimate 3000 UHPLC coupled to an Orbitrap Eclipse Tribrid mass spectrometer (Thermo Fisher Scientific). The peptides were firstly loaded onto a 75 μm × 2 cm precolumn and separated on a C18 analytical column (Acclaim PepMap 100; 75 µm × 150 mm, 3 µm particle size; Thermo Fisher Scientific) with a binary buffer system: Buffer A (0.1% formic acid in 100% H_2_O), and buffer B (0.1% FA in 80% acetonitrile with 20% H_2_O). Briefly a linear gradient of 0-40% buffer B was used to separate the peptide. The Eclipse mass spectrometer was operated in positive ion mode using data-dependent acquisition with a 3 s cycle time. Precursors and products were detected in the Orbitrap analyser at a resolving power of 60,000 and 30,000 (@ m/z 200), respectively. Raw data file was processed using Proteome Discover 3.0 (Thermo Fisher Scientific), and glycan was analysed using Byos (Protein Metrics).

### Molecular dynamics simulations

#### System Preparation and Setup

Three independent molecular systems were constructed utilising the Membrane Builder module on the CHARMM-GUI^68^ platform: (1) an isolated control lipid bilayer, (2) a monomer-embedded membrane complex, and (3) a dimer-embedded membrane complex. For the protein-containing assemblies, the respective monomeric and dimeric atomic coordinates derived from cryo-EM structural determinations were inserted into a complex lipid bilayer. The host membrane was aligned perpendicularly to the *Z-*axis and composed of a symmetric mixture of Cholesterol, POPC, POPE, and POPS in a 20:50:20:10 molar ratio. Taking the dimeric assembly as the largest representative simulation volume, the complex was explicitly solvated with 269,224 TIP3P water molecules within a rectangular periodic boundary box measuring approximately 289.6 x 289.6 x 136.1 Å^3^. Bulk ionic concentrations were modulated to 0.15 M by adding K^+^ and Cl^-^ counterions to mimic a physiological environment while achieving overall charge neutrality.

System equilibration followed a rigorous multi-stage protocol. Energy minimisation was executed in two consecutive stages: initially applying rigid harmonic position restraints to all protein atoms, followed by a completely unrestrained relaxation step. Initial equilibration runs were conducted under the standard CHARMM-GUI multi-step equilibration protocol, which features a series of short molecular dynamics intervals that systematically relax the harmonic force constraints applied to the protein coordinates and lipid headgroups. Following this relaxation phase, long unrestrained production trajectories were collected under the NPT ensemble, tracking multi-microsecond timelines for the dimeric and monomeric systems to observe equilibrium transformations and lipid translocation metrics.

#### Simulation Parameters

All explicit-solvent molecular dynamics trajectories were simulated utilising the GROMACS^69^ package driven by the CHARMM36m force field parameters^70^. Equations of motion were integrated via a standard leap-frog algorithm utilising an integration time step of 2 fs. Constant temperature was maintained at 300 K across the system via coupling to a Nosé-Hoover thermostat^71^. Continuous barostatting was enforced at an equilibrium pressure of 1 bar using a semi-isotropic Parrinello-Rahman coupling scheme^72^, allowing the area of the lateral membrane plane (*XY*) to adjust dynamically and independently from the perpendicular membrane axis (*Z*).

Long-range electrostatic topologies were evaluated using the Particle Mesh Ewald (PME) summation technique^73^ with an explicit real-space grid cutoff distance of 1.2 nm. Van der Waals forces were captured using a force-switching function operating smoothly across a distance of 1.0 to 1.2 nm. Intramolecular bond geometries for water molecules were rigidly maintained utilising the SETTLE approach^74^, whereas covalent bonds within the protein and lipid populations were regulated using the Parallel Linear Constraint Solver (P-LINCS) algorithm^75^.

#### Analysis

All post-processing spatial calculations and coordinate tracking were conducted utilising the MDAnalysis library^76^ coupled with custom Python mapping scripts.

2D Spatial Thickness Mapping: To measure localised bilayer thinning, a 2D coordinate grid was mapped onto the horizontal *XY* plane centred around the protein’s centre of mass (or the box centre for the control bilayer) with a uniform grid bin size of 2.0 x 2.0 Å^2^. For every trajectory frame, the local membrane thickness was quantified by selecting the phosphorus (P) atoms of the POPC, POPE, and POPS headgroups and tracking the absolute vertical distance between the upper and lower leaflets (Z_upper_ - Z_lower_). Time-averaged matrix cells were subsequently normalised into a thickness deviation profile (ΔThickness) by subtracting a constant baseline reference value (43.78 Å), which was computed as the global average equilibrium thickness of the unperturbed isolated control membrane.

2D Spatial Midplane Bending Mapping: To separate macroscopic membrane bending from pure vertical leaflet compression, the vertical coordinate of the mathematical bilayer midplane was computed at each grid cell using the average leaflet position formula, (Z_upper_ + Z_lower_)/2. The overall baseline midplane value was dynamically determined by averaging the midplane coordinates of unperturbed bulk lipid cells showing valid thickness profiles. Bending deviations (ΔZ) from this zero-baseline reference were calculated over the 2.0 Å spatial grid and plotted using a symmetric diverging colormap to track directional upward (positive, *+Z*) and downward (negative, *-Z*) out-of-plane warping.

## Statistical analysis

Statistical analyses were performed using GraphPad Prism 4 program (GraphPad Software Inc). NanoBiT interaction assays data are shown as mean ± SD from independent biological replicates. sbCTL1-lbCTL1 (N=6), sbCTL1-CTL1lb (N=6), CTL1sb-lbCTL1 (N=6) and CTL1sb-CTL1lb (N=6). CTL1sb + lbCTL1 (N=5), CTL1sb + lbCTL1 474 (N=5), CTL1sb + lbCTL1 477 (N=5), CTL1sb + lbCTL1 519 (N=4), CTL1sb 474 + lbCTL1 (N=5), CTL1sb 477 + lbCTL1 (N=4), CTL1sb 519 + lbCTL1 (N=3), CTL1sb 474 + CTL1lb 474 (N=4), CTL1sb 474 + CTL1lb 477 (N=4), CTL1sb 477 + CTL1lb 474 (N=4), CTL1sb 477 + CTL1lb 477 (N=5), CTL1sb 519 + lbCTL1 519 (N=5). Differences between experimental groups were evaluated by two-way ANOVA to assess the effects of each experimental factor and their interaction. When appropriate, Dunnett’s multiple-comparisons test was applied to compare each mutant condition with the corresponding wild-type control. Statistical significance was defined as *p* < 0.05.

Phospholipid thermostability assays were performed as independent experiments (N=9) and analysed separately. Differences in phospholipid-mediated stabilisation between wild-type and mutant CTL1 were assessed using two-tailed unpaired Welch’s *t*-tests. Statistical significance was defined as *p* < 0.05.

Statistical Ensemble Deformation: To compare population-wide thickness variations across the three structural configurations, the mean local membrane thickness deformation was statistically aggregated across the full production trajectories. The arithmetic means along with their corresponding standard errors of the mean (SEM) were evaluated by partitioning the data into independent blocks (N = 426 for the control, N = 426 for the monomer, and N = 418 for the dimer) to account for time-correlation (autocorrelation). A Two-Sample Welch’s *t*-test was used to determine the statistical significance of the distinct deformation gap between the monomeric and dimeric states.

## Data availability

The cryo-EM coordinates and associated maps of monomeric and dimeric CTL1 have been deposited in the Electron Microscopy Data Bank (EMDB) under accession numbers EMD-58636 and EMD-58637, respectively. Each deposition includes a corresponding full map and both half-maps. The corresponding atomic coordinates for monomeric and dimeric CTL1 have been deposited in the Protein Data Bank (PDB) under accession numbers 31RB and 31RC, respectively. Molecular dynamics simulation setup files, topology configurations, and trajectory coordinate datasets are deposited in the Molecular Dynamics DataBank (MDDB) and are available at https://mdposit.mddbr.eu/#/browse?search=redefinition%20of%20CTL1. Data that support this study are provided with this paper.

## Acknowledgments

The authors thank Israel S. Fernández for assistance in cryo-EM specimen preparation. This research has been supported by grants PID2022-137310NB-I00 (to E.E.-M.), PID2023-146771NA-I00 (to I.T.), PID2022-143177NB-I00 (to I.U.-B.), PID2021-122478NB-I00 and PCI2022-134976-2 (to M.O.) all funded by Spanish Ministry of Science, Innovation and Universities, Spanish Research Agency, and European Regional Development Fund (MICIU/AEI/10.13039/501100011033/FEDER, UE). Research in the C.V.R. laboratory is supported by the European Research Council (ERC) Advanced Grant (EP/Y029259/1). H.J. acknowledges financial support received by the UPV/EHU under the call for the recruitment of predoctoral research staff in the UPV/EHU 2024, PIF24/133. K.A.H. and M.O. acknowledge the EuroHPC project ID EHPC-EXT-2024E01-045) executed at the Barcelona Supercomputing Center. High-resolution cryo-EM data collection was performed at the Basque Resource for Electron Microscopy supported primarily by the Department of Education and the Innovation Fund of the Basque Government, the Fundación Biofísica Bizkaia, and the Spanish Ministry of Science and Innovation, through the Plan de Recuperación, Transformación y Resiliencia (PRTR) funded by the NextGenerationEU (PRTR-C17.I1) program.

## Author contributions

I.T. and E.E.-M. conceived the project. B.O.-L. and E.E.-M. generated expression constructs. A.M.-J. and E.E.-M. performed cell-based and liposome-reconstituted functional assays. A.M.-J. and G.P.-E. performed NanoBiT dimerisation assays. H.J. prepared the samples for cryo-EM. H.J. and J.P.L.-A. collected cryo-EM data. H.J. and I.T. proceeded with cryo-EM data processing, model building and structure refinement. K.A.H and M.O. designed and carried out the molecular dynamics-related study. H.S. and C.V.R performed and analysed the native mass spectrometry, lipidomics and proteomics experiments. H.J. wrote the original draft. H.J., I.U.-B, I.T. and E.E.-M edited the manuscript. All authors read and approved the final manuscript.

## Supplementary information

Supplementary information includes Supplementary Figures 1-14, Supplementary Movies 1-3, and Supplementary Tables 1-2.

## Materials & Correspondence

Correspondence and requests of materials should be addressed to Iban Ubarretxena-Belandia, Igor Tascón or Ekaitz Errasti-Murugarren

