## Supplementary figures for "Structural basis of selective phospholipid recognition and local membrane deformation by CTL1"

Hanxing Jiang *et al.*

*Corresponding authors: (E.E-M), (I.T), and (I.U-B)

**This Supplementary Information includes:**

Supplementary Figures 1 to 14

Supplementary Video Legends 1 to 3

Supplementary Tables 1 to 2

Supplementary References

**Other Supplementary Materials for this manuscript include the following:**

Supplementary Videos 1 to 3 (mp4 files)

Supplementary Table 1 (Excel file)

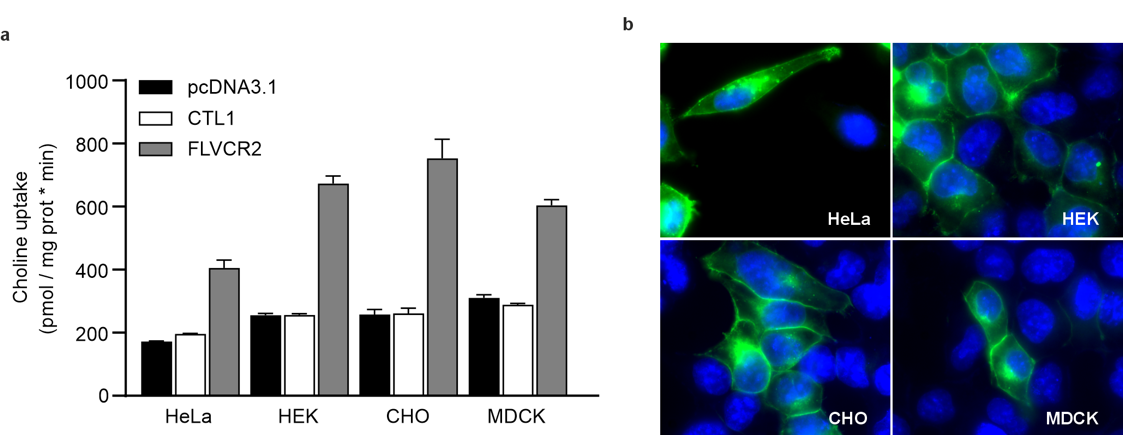

**Supplementary Figure 1. Comparison of FLVCR2- and CTL1-mediated choline uptake in different cell lines. a** Initial rates of [^3^H]-choline uptake (pmol choline/mg protein•min) in HeLa, HEK293T, CHO and MCDK cells transiently expressing human CTL1 (CTL1-pcDNA3.1), human FLVCR2 (FLVCR2-pcDNA3.1), or empty vector (pcDNA3.1). Uptake was measured after a 1 min incubation at 10 μM choline, in Na^+^-containing (137 mM NaCl) transport buffers at extracellular pH 7.5. CTL1 overexpression did not increase choline uptake above control cells in any of the assayed cell lines, whereas FLVCR2 consistently mediated robust choline uptake. Data (mean ± SD) correspond to three independent experiments run in quadruplicate (N=3). **b** Subcellular localisation of GFP-tagged human CTL1 in HeLa, HEK293T, CHO and MDCK cells. Colour code: CTL1-GFP (green) and DAPI (blue).

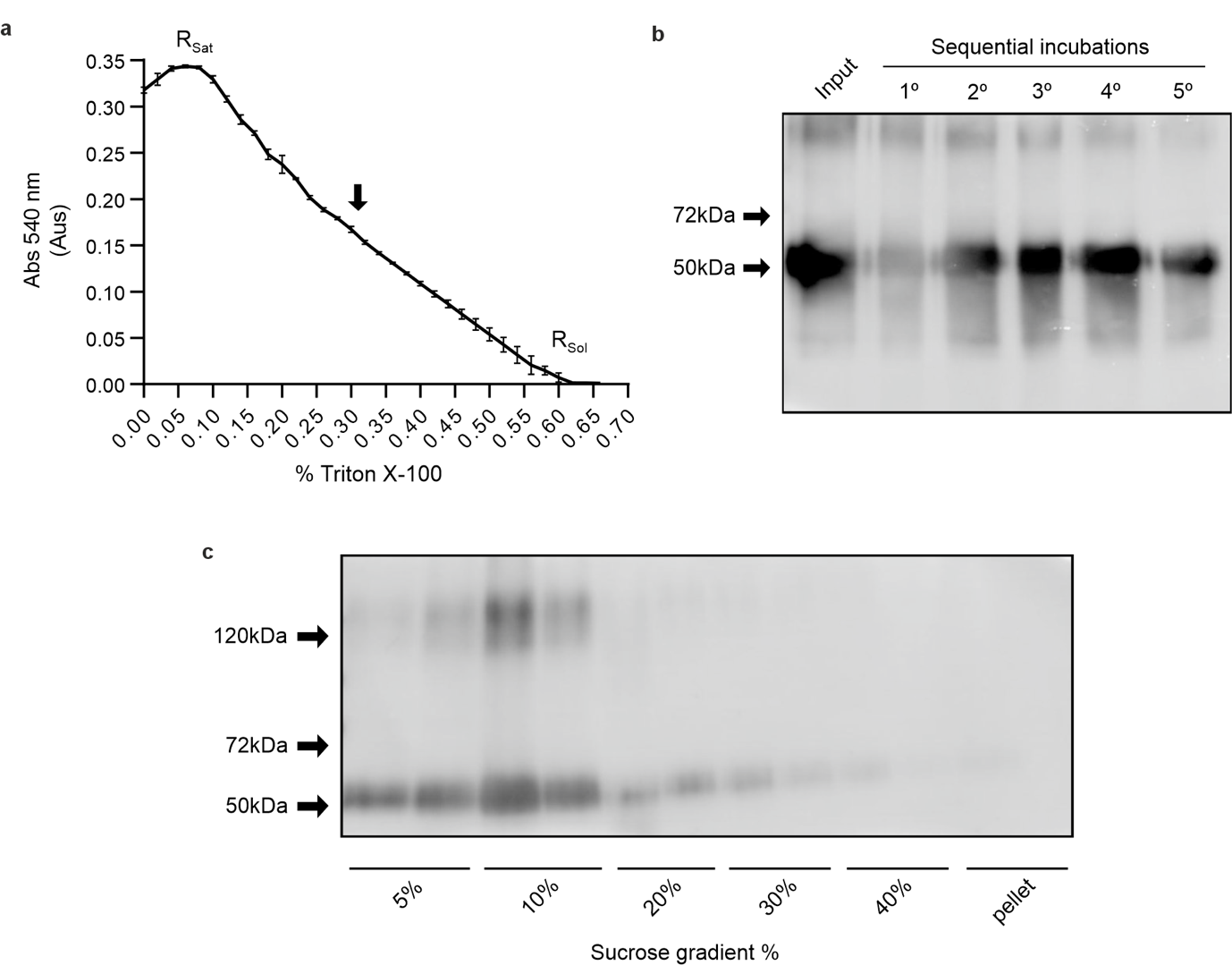

**Supplementary Figure 2. CTL1 reconstitution in total brain lipid extract liposomes. a** Determination of the detergent saturation (R_sat_) and solubilisation (R_sol_) ratios for total brain lipids liposomes in the presence of Triton X-100. The middle arrow indicates the point in the titration curve at which the liposomes were combined with purified CTL1. **b** Western blot analysis of protein remaining in solution during detergent removal by sequential Bio-Beads incubations. Lane 1 corresponds to the sample before the first Bio-Beads incubation, followed by samples collected after each of the five successive incubations. **c** Sucrose density gradient fractionation of the reconstituted proteoliposomes.

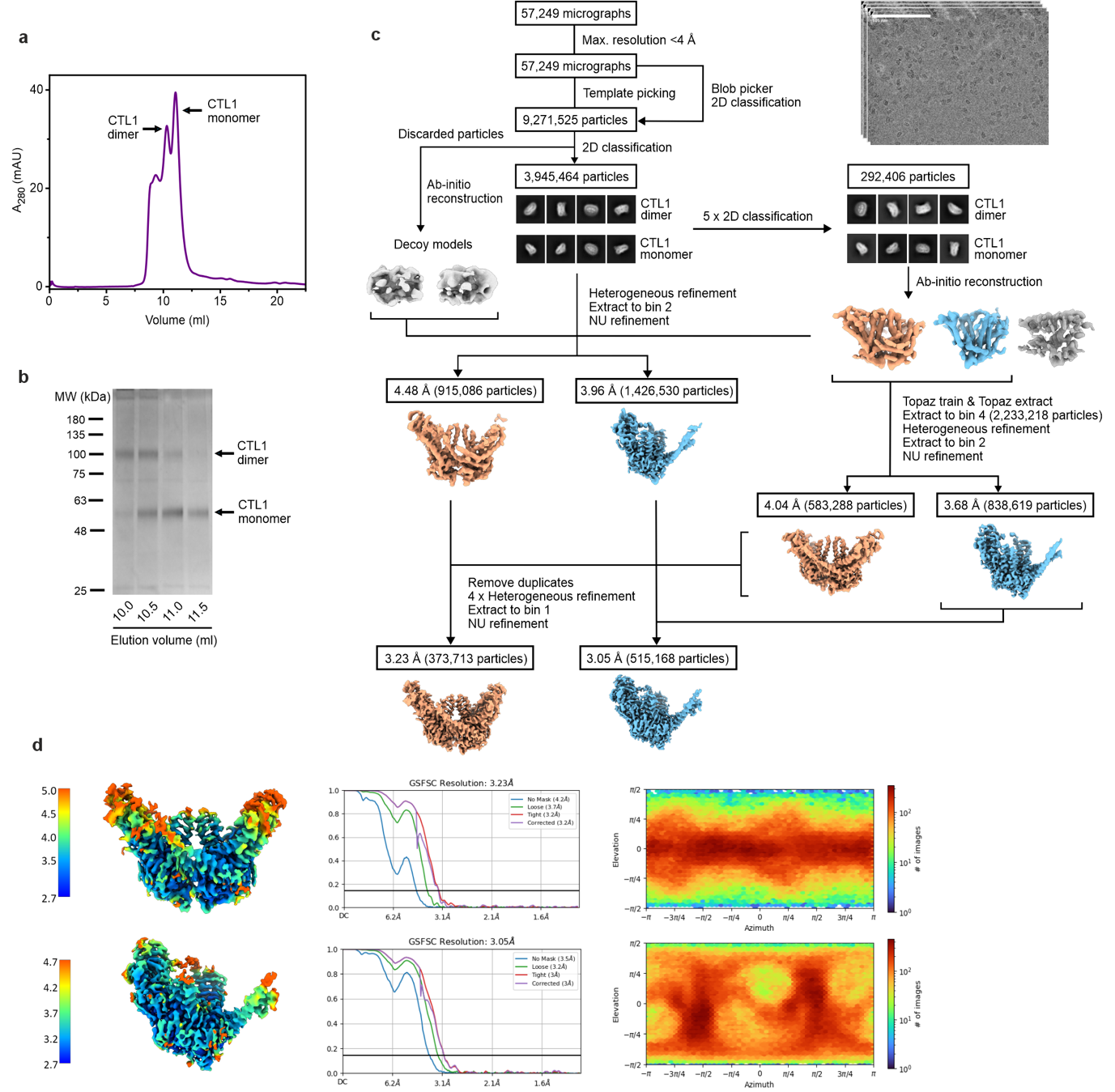

**Supplementary Figure 3. Purification and cryo-EM structure determination of human CTL1. a** Size-exclusion chromatography profile of a representative purification of human CTL1. **b** SDS-PAGE showing bands corresponding to purified CTL1 after size-exclusion chromatography. **c** Cryo-EM data processing diagram of CTL1. **d** Dimeric (top) and monomeric (bottom) CTL1 coloured by local resolution as estimated using CryoSPARC (left), gold-standard FSC curves (centre) and Euler diagrams showing particle orientation distribution (right).

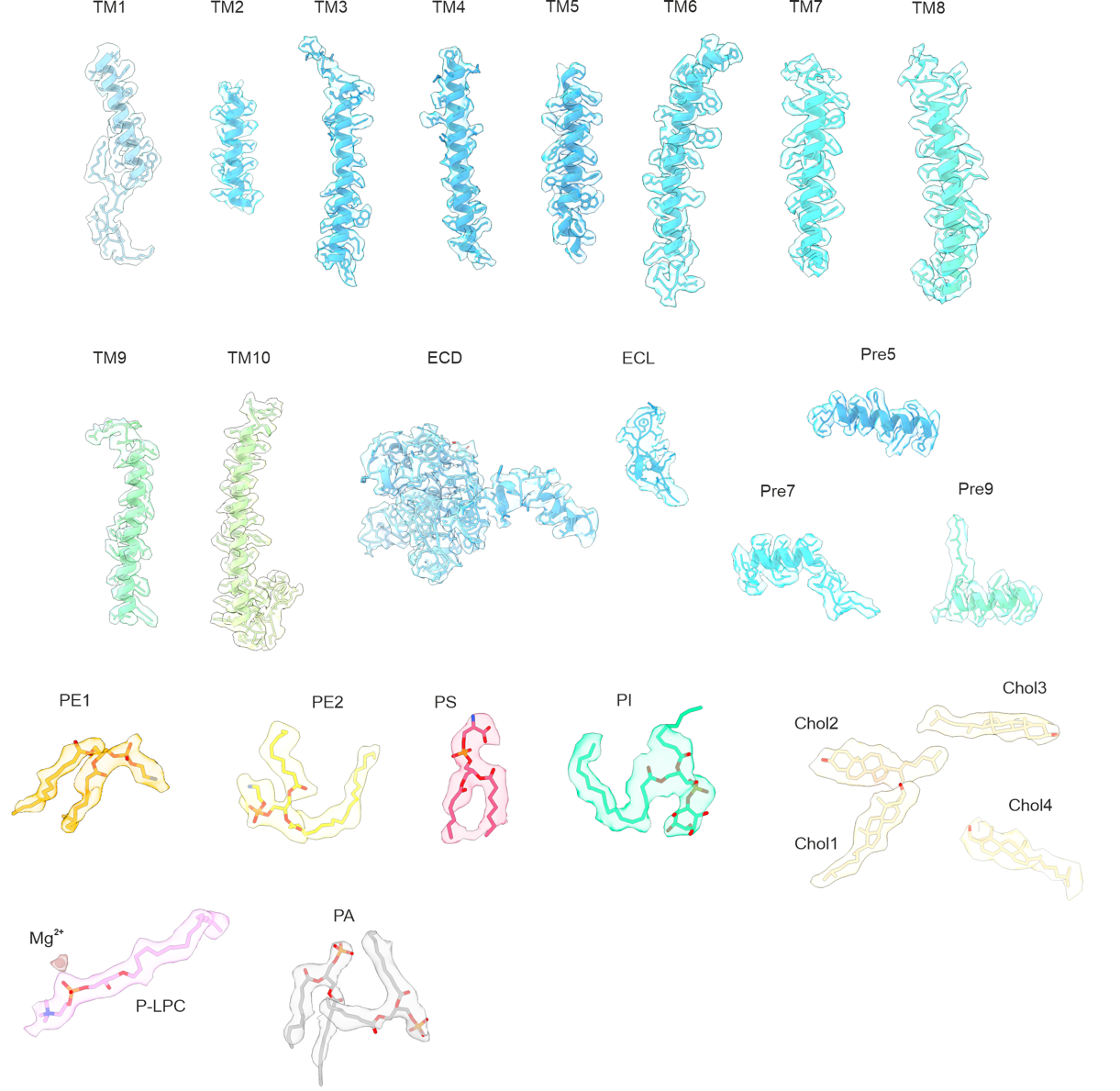

**Supplementary Figure 4. Selected cryo-EM map density regions and model-to-map fit of CTL1.** Cartoon representation of different segments of CTL1 and associated ligands within the cryo-EM densities, coloured as in **Figure 2**. Side chains and ligands are shown as sticks.

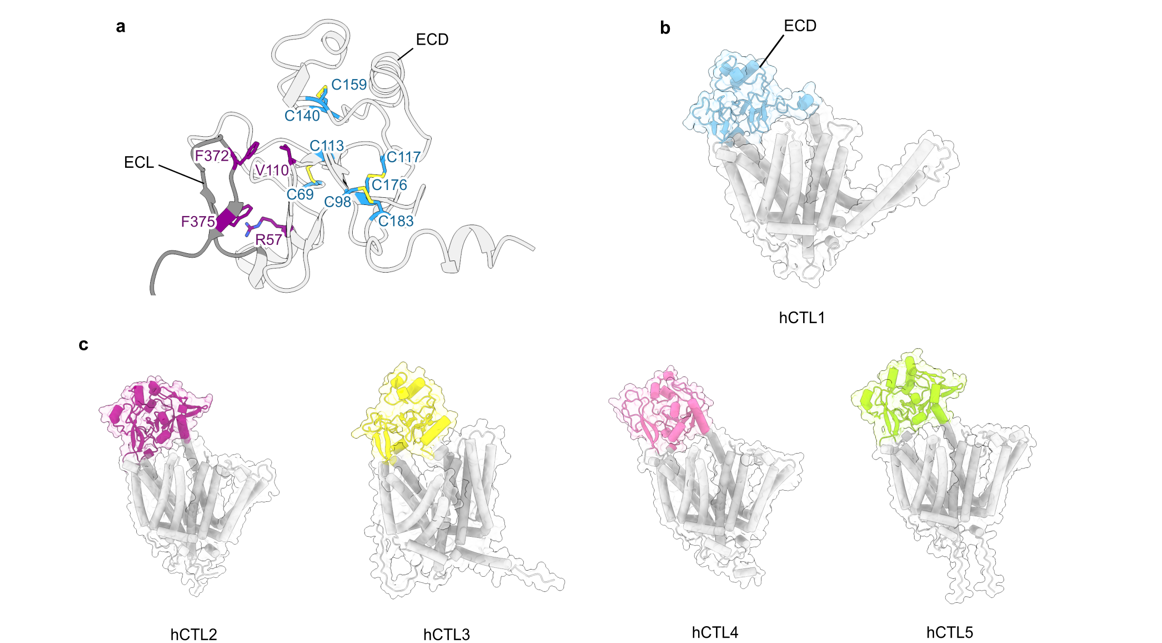

**Supplementary Figure 5. Comparison of the CTL1 ECD with AlphaFold3 predictions for CTL2-5. a** Cartoon representation of the CTL1 ECD (light grey) showing cystine pairs (blue sticks) that stabilise the overall domain, and residues mediating interactions between the ECL (dark grey) and ECD are shown as purple sticks. **b** Cartoon representation of experimental CTL1 structure with the ECD depicted in light blue and the TM helices in white. **c** AlphaFold3^1^-predicted structures of CTL2-5 with the ECD highlighted in different colours.

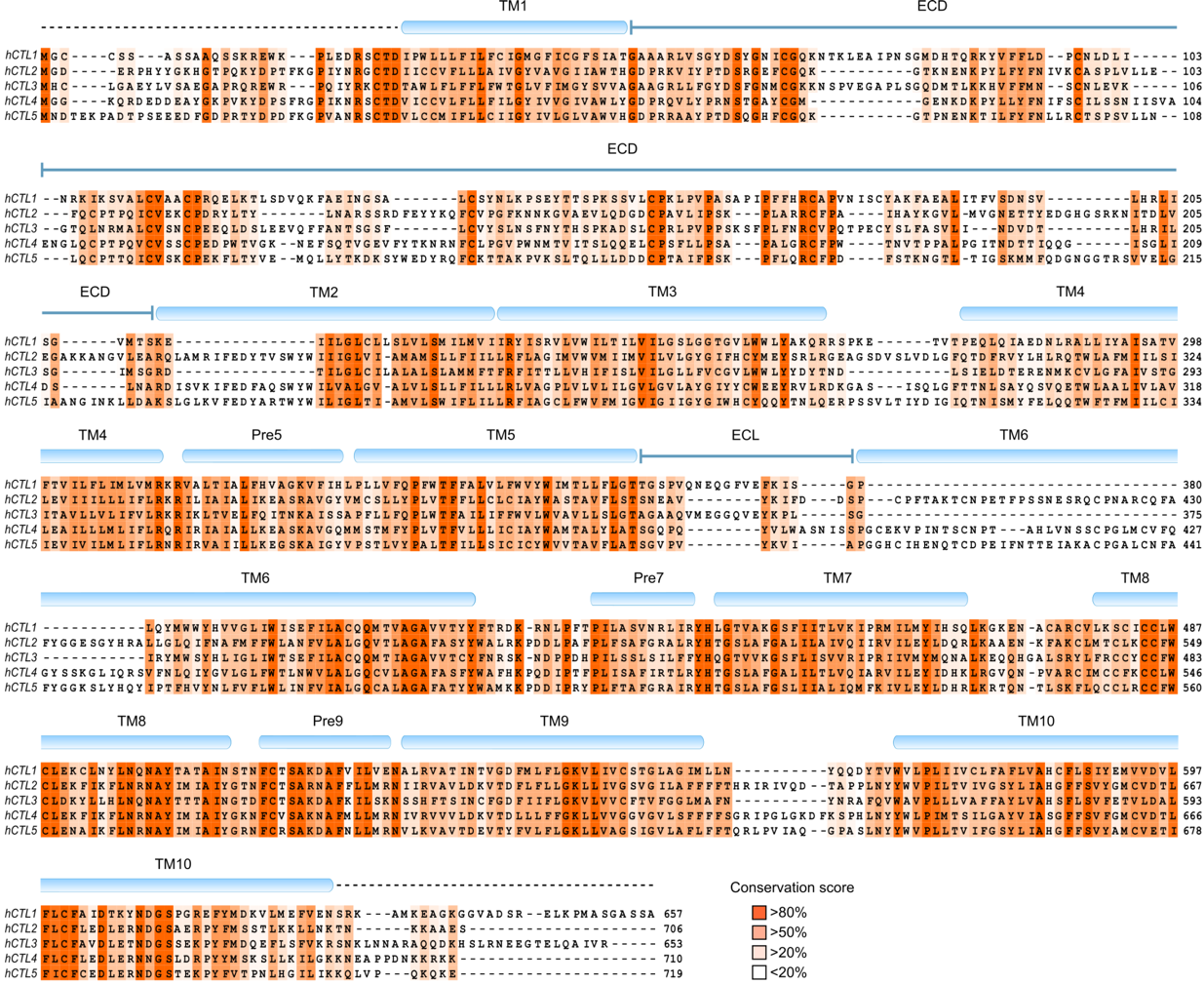

**Supplementary Figure 6. Sequence alignment of human CTL1-5.** Residues are coloured according to sequence conservation. Secondary motifs of CTL1 are depicted as light blue cylinders, and dashed lines at N- and C-termini indicate unresolved residues in the CTL1 cryo-EM structure. The sequence alignment was carried out using PROMAL3D and visualised with Jalview.

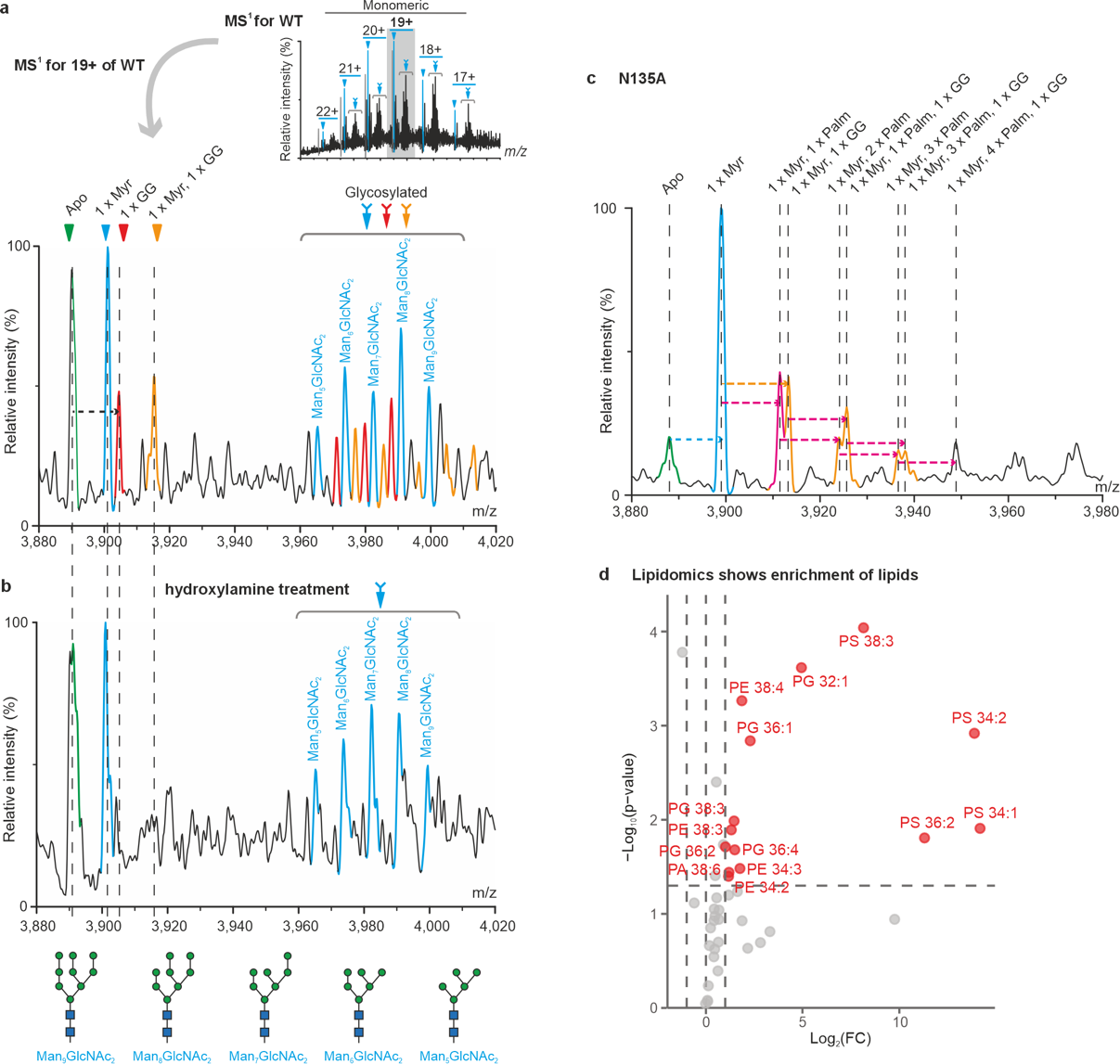

**Supplementary Figure 7. nMS analysis of CTL1 and CTL1-N135A. a** Expanded view of the 19+ charge state of wild-type monomeric CTL1. CTL1 was extensively modified by myristoylation (Myr, cyan), geranylgeranylation (GG, red), and glycosylation. **b** Expanded view of 19+ charge state of monomeric CTL1 following hydroxylamine treatment. Native mass spectrum shows reduced palmitoylation and geranylgeranylation. **c** Expanded view of the 19+ charge state of monomeric N135A, which clearly shows extensive post-translational modifications. **d** Volcano plot of lipids enriched with purified wild-type CTL1. Phospholipids identified as enriched in CTL1 samples are shown, including PA, PE, PG, PI, and PS. The x-axis shows log_2_ fold change relative to flow-through fractions, and the y-axis shows −log_10_(p-value).

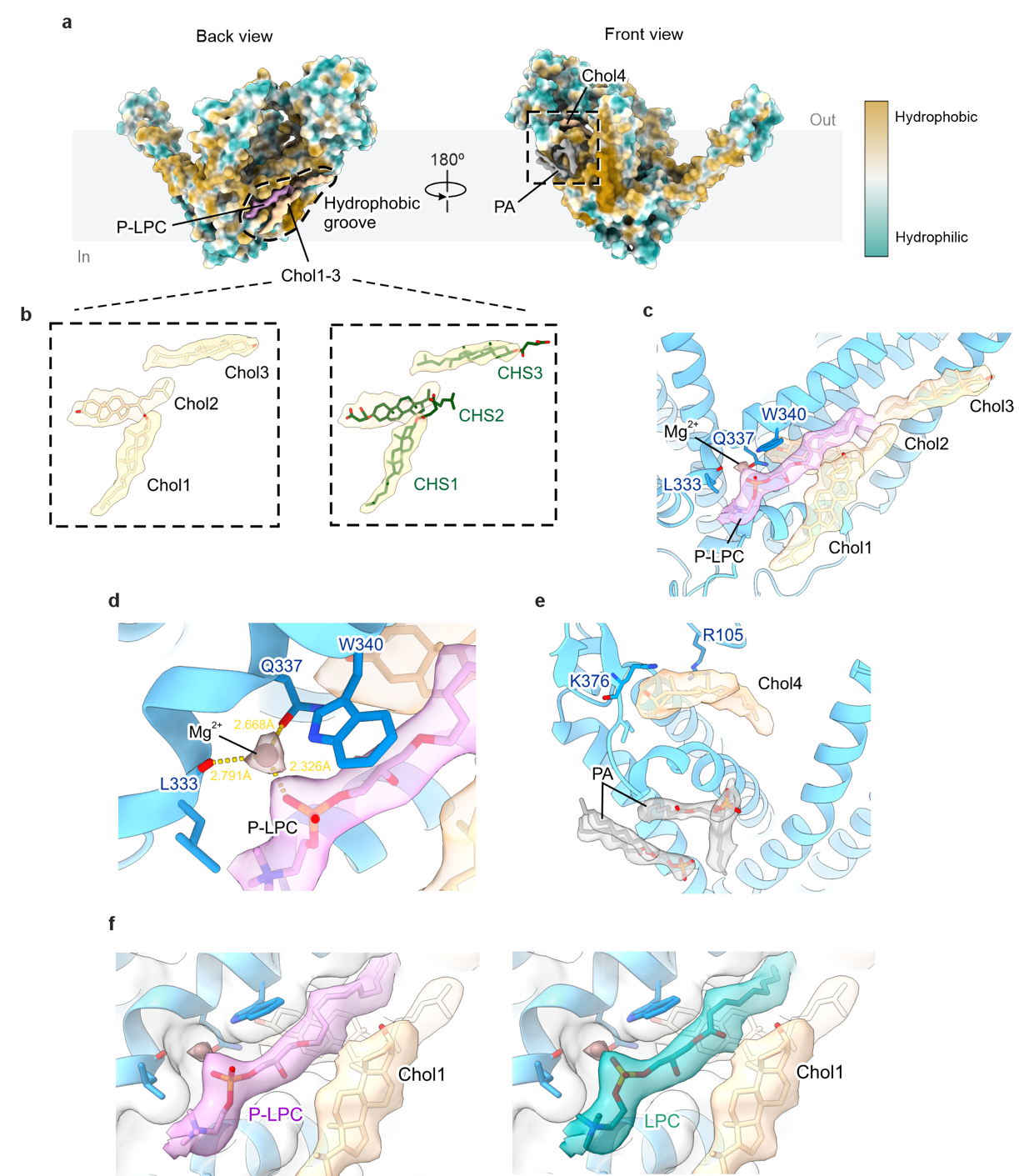

**Supplementary Figure 8. Lipid interactions within CTL1 hydrophobic groove. a** Surface representation of CTL1 coloured by hydrophobicity. Left, back view shows cholesterols (Chol1-3), plasmalogen lysophosphatidylcholine (P-LPC) and Mg^2+^ ion densities within the hydrophobic groove indicated by dashes lines. Right, dashed box in the front view shows PA and Chol4 densities attached to CTL1. **b** Comparison of cholesterol (left; beige) and CHS (right; green) molecules fitted into the cholesterol-like cryo-EM densities in the hydrophobic groove. **c** Close-up view of the cartoon representation of the hydrophobic groove (blue), outlined in **a (left)**, showing the cluster of three cholesterol molecules (Chol1-3; light beige) and a P-LPC molecule (light pink) fitted into their respective densities. Mg^2+^ ion is shown as brown sphere and closely interacting residues as blue sticks. **d** Close-up view of the Mg^2+^ ion coordination with surrounding residues and P-LPC. **e** Close-up view of the binding region of PA (grey) and Chol4 (light beige) molecules within the corresponding densities as outlined in **a (right)**, with Chol4-interacting residues as blue sticks. **f** Comparison of P-LPC (left; light pink) and lysophosphatidylcholine (LPC; right, teal) molecules fitted into the elongated cryo-EM density within the hydrophobic groove. The vinyl ether moiety of P-LPC allows hydrophobic contacts with the sterol rings of Chol1, whereas in LPC the corresponding position is occupied by an ester carbonyl group.

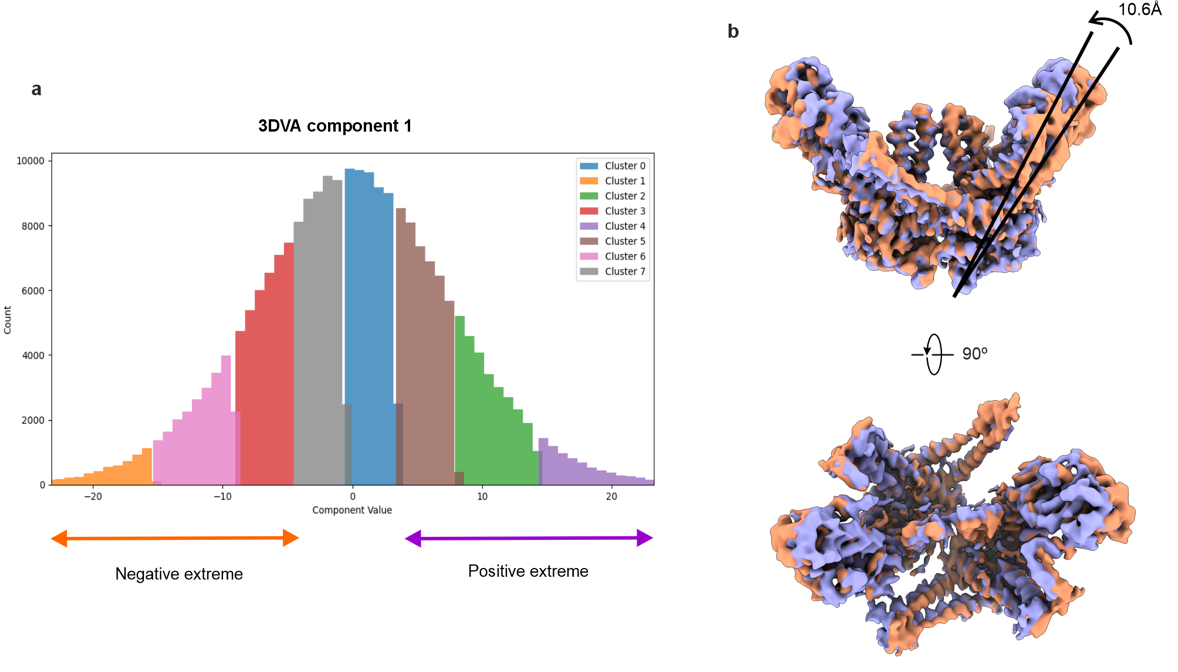

**Supplementary Figure 9. 3D variability analysis of dimeric CTL1. a** Particle distribution histogram of the first variability component obtained in CryoSparc. **b** Density maps corresponding to the negative (orange) and positive (purple) extremes of the first component are shown, revealing continuous motion of the ECD relative to the core region. Black lines indicate the direction and amplitud of the motion (10.6 Å).

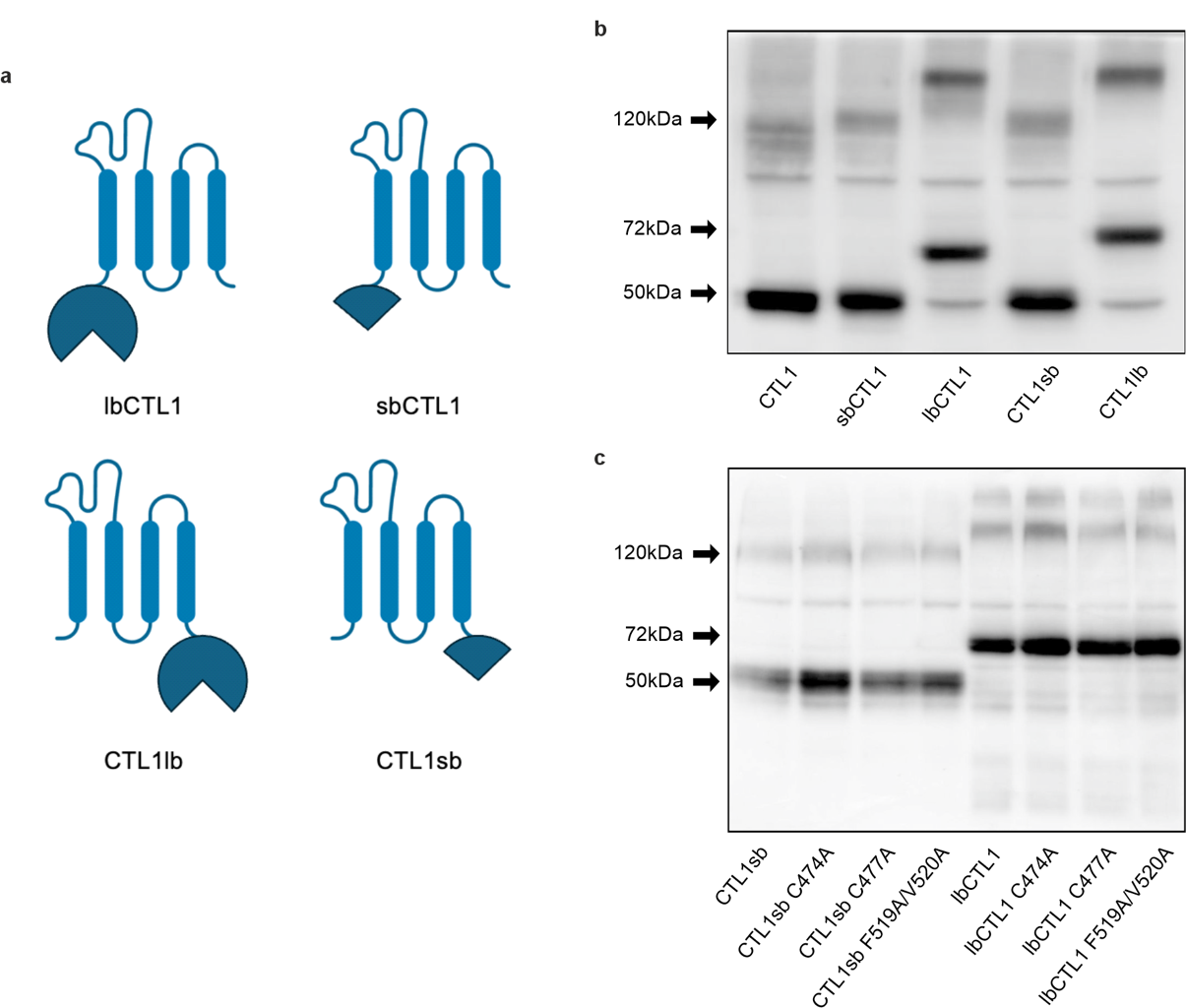

**Supplementary Figure 10. Expression levels of Small BiT (sb) or Large BiT (lb) linked wild-type CTL1 and variants. a** Schematic depiction of SmBiT/LgBiT-tagged CTL1 constructs to identify the optimal combination for the NanoBiT assay. Western blot analysis of wild-type (**b**) and CTL1 variants (**c**) linked to either Small BiT (sb) or Large BiT (lb). Pelleted HEK293T cells overexpressing the corresponding construct were lysed and aliquots resolved by 10% SDS-PAGE and blotted onto nitrocellulose membranes. Protein detection utilised anti-CTL1 antibody (14687-1-AP) and was performed by enhanced chemiluminescence, using as a secondary goat anti-rabbit antibody conjugated to horseradish peroxidase.

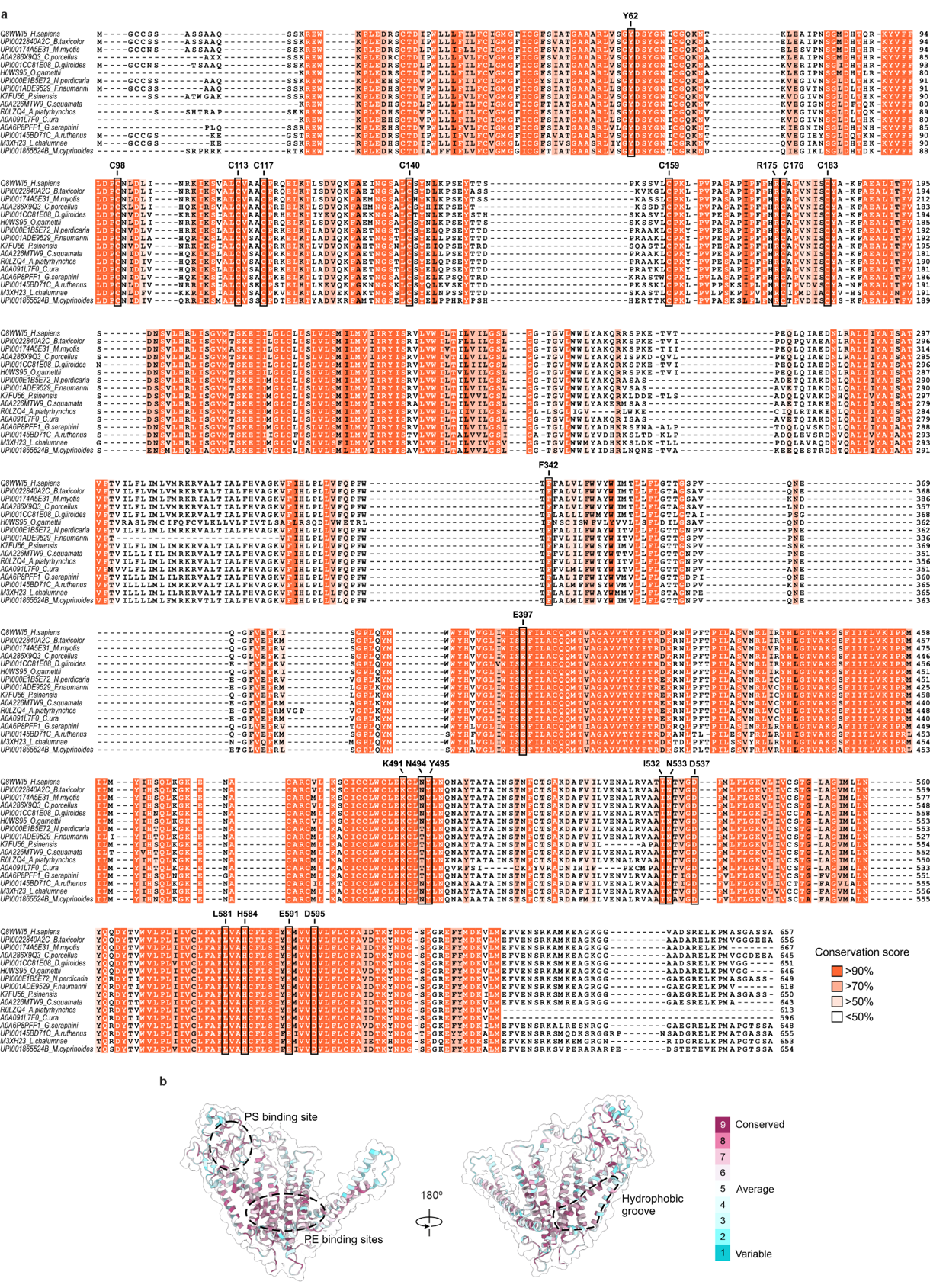

**Supplementary Figure 11. Sequence conservation analysis of CTL1 homologs. a** Top 15 most similar CTL1 homologs among 150 sequences identified by ConSurf. Residues are coloured based on sequence conservation, and residues of interest are annotated. **b** Conservation mapping of CTL1 in front (left) and back (right) views, with residues coloured based on the conservation scores. Dashed lines indicate highly conserved regions corresponding to the PS-binding site, PE-binding sites, and hydrophobic groove.

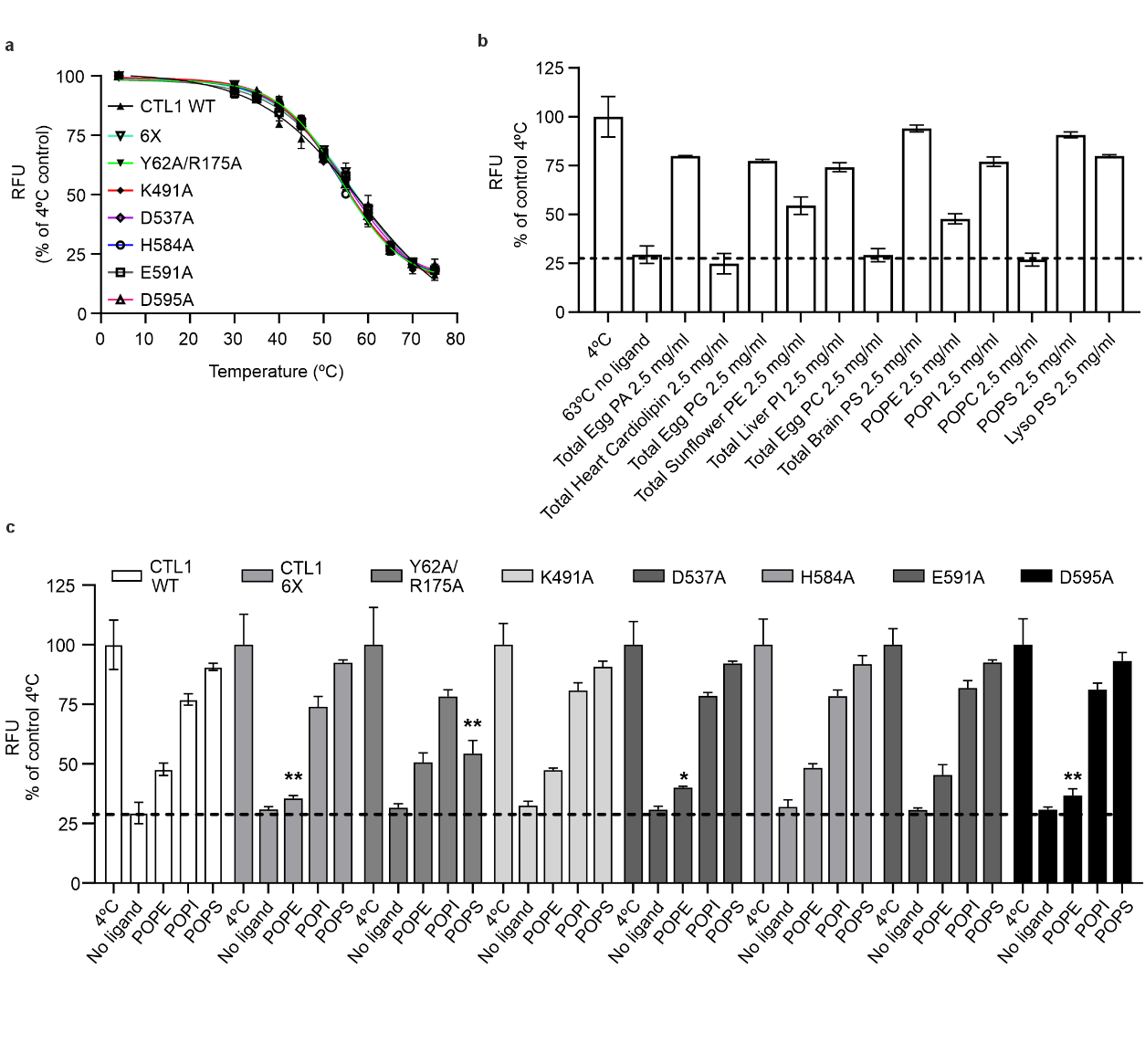

**Supplementary Figure 12. Phospholipid-dependent stabilisation of CTL1 is mediated by distinct lipid-binding residues. a** Thermal stability of NG-delipidated wild-type CTL1-GFP and the indicated phospholipid-binding mutants was assessed using a GFP-based thermal shift assay. Fluorescence was normalised to the signal at 4 °C and fitted to a sigmoidal dose-response curve to determine the melting temperature (*Tm*). The *Tm* values were 57.51 ± 0.64 °C (WT), 53.95 ± 0.17 °C (6X), 54.36 ± 1.39 °C (Y62A/R175A), 55.26 ± 0.86 °C (K491A), 53.77 ± 0.94 °C (D537A), 56.26 ± 0.59 °C (H584A), 54.34 ± 1.93 °C (E591A) and 56.60 ± 0.48 °C (D595A). Data are presented as mean ± SD from three independent protein purifications run in triplicate (N=3). **b** Effect of phospholipids on CTL1 thermal stability assessed by GFP-based thermal shift assay. Wild-type CTL1-GFP was incubated with the indicated phospholipids (2.5 mg/ml) or vehicle before heat treatment at 63°C (5°C above the *T_m_* of wild-type CTL1). Fluorescence remaining after heat treatment was normalised to the fluorescence measured at 4 °C. Data are presented as mean ± SD from three independent protein purifications run in triplicate (N=3). **c** Effect of phospholipids on the thermal stability of wild-type CTL1-GFP and phospholipid-binding mutants. Purified proteins were incubated with the indicated phospholipids (2.5 mg/ml) before heat treatment at a temperature 5°C above the corresponding *T_m_* of each construct. Fluorescence remaining after heat treatment was normalised to the fluorescence measured at 4 °C. Mutations 6X, D537A and D595A significantly reduced POPE-mediated stabilisation, whereas the Y62A/R175A mutant displayed reduced stabilisation by POPS. Data are presented as mean ± SD from three independent protein purifications run in triplicate (N=3). Statistical significance was determined using two-tailed unpaired Welch's *t*-tests. *p* < 0.05 (*), *p* < 0.01 (**). POPE WT vs 6X: *p*: 0.0062, *t* value: 7.44 and *df*: 2.815; POPE WT vs D537A: *p*: 0.0298, *t* value: 5.063 and *df*: 2.212; POPE WT vs D595A: *p*: 0.0074, *t* value: 5.032 and *df*: 3.978; POPS WT vs Y62A/R175A: *p*: 0.0049; *t* value: 11.06 and *df*: 2.815.

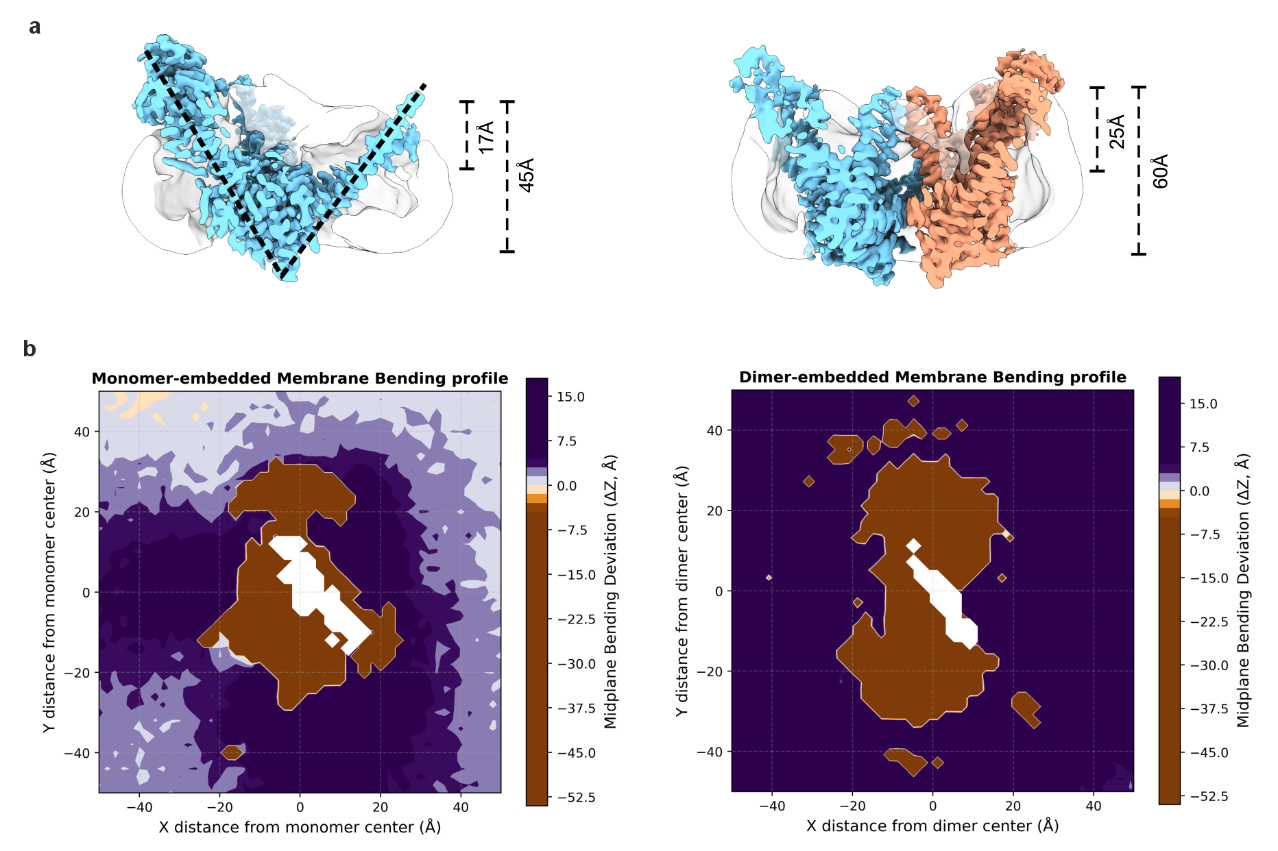

**Supplementary Figure 13. Membrane bending deviation induced by monomeric and dimeric CTL1. a** Cross-sectional view of the cryo-EM maps of monomeric (left) and dimeric (right) CTL1 showing detergent micelles (white) narrow toward the central cavity. The dashed line in the monomer map highlights its conical shape. **b** 2D spatial profiles of local bilayer midplane bending for monomer-embedded (left), and dimer-embedded membrane (right). Overall bilayer bending was calculated as the time-averaged vertical position of the membrane midplane computed as (Z_upper_+Z_lower_)/2 on a 2.0 Å XY grid centred around the protein centre of mass. Bending deviations (ΔZ) were normalised relative to the average unperturbed membrane midplane position, with ΔZ = 0 Å indicating a flat membrane. Positive values (purple regions) indicate local upward buckling toward the extracellular space, whereas negative values (orange regions) denote downward depression toward the intracellular space. Central masked patches trace the footprint of the protein core.

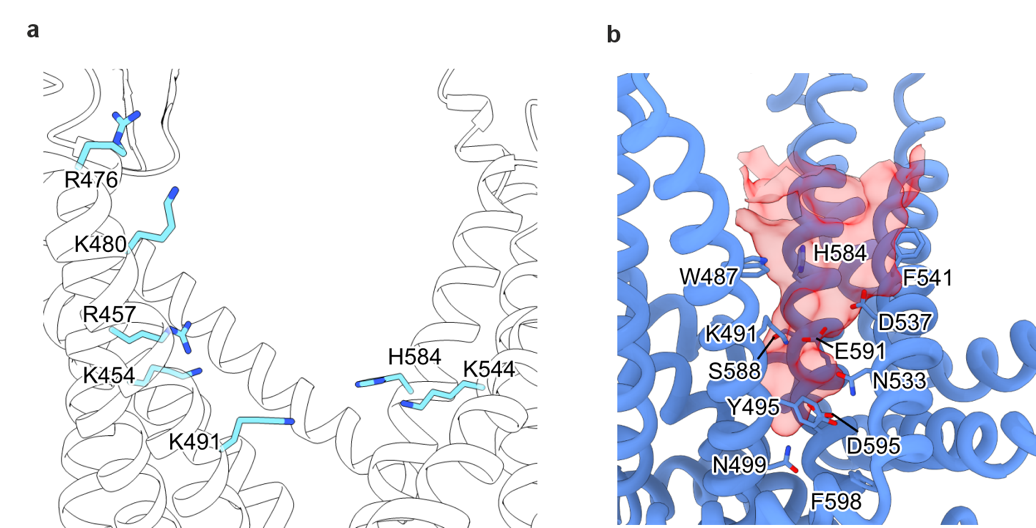

**Supplementary Figure 14. Water-accessible pathway from extracellular side toward the central cavity of CTL1**. **a** Cartoon representation of the polar network region in CTL1 formed by highly conserved polar and positively charged residues (light blue sticks) extending from the extracellular vestibule toward the PE1-binding site. **b** Representative snapshot from MD simulations displaying a transient water-accessible pathway (red) in dimeric CTL1 (blue). Residues in the pathway are shown as sticks.

**Supplementary Video legends**

**Supplementary Video 1.** MD simulation showing a surface representation of dimeric CTL1 (protomers in cyan and blue) embedded in a symmetric POPC:POPE:POPS:CHOL membrane bilayer (phosphorus atoms shown as white spheres). Surrounding phospholipids molecules (beige spheres) from the extracellular leaflet are recruited toward the central cavity of CTL1.

**Supplementary Video 2.** MD simulation showing the sequential recruitment of two phospholipid molecules (beige spheres) into binding positions 1 and 2 within the central cavity of CTL1, shown in cartoon representation.

**Supplementary Video 3.** MD simulation showing the back view of dimeric CTL1 with a POPS (beige spheres) molecule entering the PS-binding pocket located in the ECD.

**Supplementary Table**

**Supplementary Table 1.** CTL1 conservation analysis by ConSurf

(separate Excel file)

**Supplementary Table 2**. Primers used to generate CTL1 variants

| Primer name | | Sequence (5' → 3'） |
| --- | --- | --- |
| Y62A | Forward | CCTGGTGTCCGGCGCCGACAGCTACGG |
|  | Reverse | CCGTAGCTGTCGGCGCCGGACACCAGG |
| N135A | Forward | CAAAAGTTCGCCGAAATCGCCGGCAGCGCCCTCTGTAG |
|  | Reverse | CTACAGAGGGCGCTGCCGGCGATTTCGGCGAACTTTTG |
| R175A | Forward | CCCTATCCCTTTTTTTCACGCCTGCGCCCCTGTGAAC |
|  | Reverse | GTTCACAGGGGCGCAGGCGTGAAAAAAAGGGATAGGG |
| C474S | Forward | CAAGGAGAACGCCTCCGCCAGATGCGTG |
|  | Reverse | CACGCATCTGGCGGAGGCGTTCTCCTTG |
| C477S | Forward | CGCCTGCGCCAGATCCGTGCTGAAGTCTTG |
|  | Reverse | CAAGACTTCAGCACGGATCTGGCGCAGGCG |
| F519A/  V520A | Forward | CCAGCGCCAAGGACGCCGCCGCCATCCTGGTGGAAAACG |
|  | Reverse | CGTTTTCCACCAGGATGGCGGCGGCGTCCTTGGCGCTGG |
| K491A | Forward | GCCTGTGGTGCCTGGAGGCCTGCCTGAACTACCTGAA |
|  | Reverse | TTCAGGTAGTTCAGGCAGGCCTCCAGGCACCACAGGC |
| D537A | Forward | CCATCAATACAGTGGGCGCCTTCATGCTGTTTCTGGG |
|  | Reverse | CCCAGAAACAGCATGAAGGCGCCCACTGTATTGATGG |
| H584A | Forward | TTTGCTTTTCTCGTGGCCGCCTGCTTCCTGTCTATCTAC |
|  | Reverse | GTAGATAGACAGGAAGCAGGCGGCCACGAGAAAAGCAAA |
| E591A | Forward | CTGCTTCCTGTCTATCTACGCCATGGTGGTAGACGTGTTA |
|  | Reverse | TAACACGTCTACCACCATGGCGTAGATAGACAGGAAGCAG |
| D595A | Forward | CTACGAGATGGTGGTAGCCGTGTTATTCCTGTGC |
|  | Reverse | GCACAGGAATAACACGGCTACCACCATCTCGTAG |
